# Unconventionally primed type 1 follicular helper T cells are required for long-lived IgA plasma cell development following mucosal viral infection

**DOI:** 10.1101/2025.01.27.635134

**Authors:** Kei Haniuda, Natalie M. Edner, Yuko Makita, Tania H. Watts, Gregory F. Wu, Thamotharampillai Dileepan, Jennifer L. Gommerman

## Abstract

Although IgA^+^ long-lived plasma cells (LLPCs) generated following mucosal viral infection provide durable protection against reinfection, little is known about their generation. Here, we show that oral RV infection induces gut-resident LLPCs that produce highly mutated protective IgA. Unlike RV-specific IgG^+^ LLPCs, IgA^+^ LLPCs were generated independently of MHCII expression by dendritic cells – rather MHCII expression by B cells was both necessary and sufficient. B cell MHCII was also sufficient to induce a unique population of T-bet^+^ follicular helper T (T_FH_1) cells which were crucial for RV-specific IgA^+^ LLPC accumulation in the gut via IFNγ- and CXCR3-dependent mechanisms. Similar to RV infection, T_FH_1 cells were required for influenza-specific IgA response. However, unlike RV infection, B cell MHCII was not sufficient to induce influenza-specific IgA^+^ LLPCs, suggesting the operation of mucosal site-specific priming mechanisms. Collectively, our data reveal that unconventionally primed T_FH_1 cells support IgA responses to mucosal viral infections.

## Introduction

Many pathogens invade the body through mucosal surfaces such as the gut and respiratory tract. At the mucosae secreted antibodies (Abs), and in particular immunoglobulin A (IgA) produced by resident plasma cells (PCs), provide protective immunity. Secreted IgA has potent neutralizing capacity due to its dimeric form and can eliminate pathogens before they replicate in the host as observed following oral vaccination against poliovirus and rotavirus (RV)^1^. However, many licensed vaccines, including those against SARS-CoV-2, do not completely protect the host against mucosal infection^2^. This limitation is likely due to their inability to induce effective and durable mucosal IgA responses. Indeed, we and others have shown that the level of mucosal IgA induced by SARS-CoV-2 vaccination is much lower than that induced by natural infection^3–7^. Moreover, the IgA response to SARS-CoV-2 vaccination, and even SARS-CoV-2 infection, is short-lived^3,8^. Thus, to develop protective vaccines against pathogens that enter through the mucosa, we must gain a better molecular understanding of the generation of mucosal IgA responses.

An effective and durable Ab response relies on the formation of long-lived PCs (LLPCs) that produce high affinity Abs. Generation of LLPC typically requires priming of CD4^+^ T cells by Zbtb46-dependent conventional/classical dendritic cells (cDCs) within secondary lymphoid tissues^9,10^, which we term here as “induction sites”. cDCs serve as specialized Ag-presenting cells (APCs) which express high levels of MHCII and costimulatory molecules^10,11^. After priming, T cells move to the boundary of T and B cell zones where they interact with cognate B cells and fully differentiate into follicular helper T (T_FH_) cells to form germinal centers (GCs). In the GC, B cells undergo secondary diversification of their B cell receptors (BCR), including class switch and affinity maturation, the latter facilitated by rounds of somatic hypermutation (SHM) of the BCR. CD40 and cytokine signals from T_FH_ cells drive affinity maturation of GC B cells which eventually differentiate into PCs. PCs then migrate to “effector sites” such as the bone marrow (BM), where a small fraction of PCs successfully access specialized survival niche microenvironments, persisting as LLPCs for months in rodents and decades in humans^2,12,13^.

Most of our knowledge on LLPC generation and maintenance comes from studying systemic IgG responses whereas information on the induction, regulation and durability of mucosal IgA responses is more limited. Most of our mechanistic understanding is derived from studies analyzing responses to host microbiota and food antigens (Ags). These IgA responses occur throughout the lifespan of the organism^1,14^ and can occur in the absence of T cells^15^, T_FH_ cells and CD40/CD40L^16,17^. It is currently unclear how pathogen-specific IgA responses develop in mucosal tissues and whether the development of these IgA responses differs from that of IgG.

Here, we investigated molecular mechanisms underlying the development of mucosal IgA^+^ PCs following mouse infection with RV, an enteric virus which induces a *de novo* durable IgA response in the gut. We found that RV infection induces a T_FH_-dependent accumulation of IgA^+^LLPCs in the small intestinal lamina propria (SILP). These IgA^+^ LLPCs exhibit a history of clonal expansion and a high level of SHM. The development of RV-specific IgA^+^ LLPCs requires a unique T-bet^+^ IFNγ^+^ (type 1) T_FH_ subset (T_FH_1), the induction of which depends on Ag-presentation by B cells but, surprisingly, not Ag-presentation by cDCs. We further identified that IFNγ from T_FH_ cells upregulates CXCR3 on IgA^+^ PCs, and that a T_FH_1-IFNγ-CXCR3 axis is required for the accumulation of IgA^+^ PCs in the SILP. Lastly, as a comparator, we investigated mucosal IgA response in the respiratory tract following influenza A virus (IAV) infection and found that, while the development of IAV-specific IgA^+^ LLPCs partially depends on T_FH_1 cells, B cell-intrinsic MHCII expression is not sufficient for inducing an IgA response to IAV, suggesting different priming requirements between the respiratory vs intestinal mucosae. Our findings provide novel insights into mechanisms of T cell-dependent IgA responses to mucosal viral infections.

## Results

### Generation of RV-specific IgG and IgA plasma cells requires T_FH_ and CD40 signaling

RV infects small intestinal epithelial cells, inducing a *de novo*, durable and protective IgA response^18–21^. To study the mechanism responsible for generating RV-specific IgA^+^ LLPCs, we employed a virulent murine RV strain, ECw, that causes acute infection in wild type mice^22^. In our vivarium, we found that oral gavage with RV results in viral shedding into the feces until 7 days post infection (dpi) (Figure S1A). The RV spike protein (VP4) has been identified as a target of protective neutralizing antibody, and VP6, the middle capsid layer protein, is highly immunogenic^19,21,23^. To characterize RV-specific B cell responses, we produced recombinant RV VP4 and VP6 proteins (Figure S1B) and generated Ag-tetramer molecules by mixing site-specific biotinylated VP4 and VP6 with fluorochrome-conjugated streptavidin (SA). We first used these tetramers to characterize RV-specific B cells by flow cytometric analysis. In the Peyer’s patches (PPs) and gut-draining mesenteric lymph node (MLN), the induction sites for immune responses in the small intestine^14^, we detected VP4- and VP6-specific B220^+^GL7^+^CD38^-^IgD^-^IgG^+^ and B220^+^GL7^+^CD38^-^IgD^-^IgA^+^ GC B cells peaking at 14-28 dpi. These were maintained at reduced levels until at least 56 dpi (Figure 1A, 1B, S1C and S1D). We also detected VP4- and VP6-specific CD138^+^B220^low^IgG^+^ and CD138^+^B220^low^IgA^+^ cells in the PPs and MLN. These are likely a combination of plasmablasts and PCs, but for simplicity we call this collective population PCs. IgA^+^ PCs appeared in the PPs and MLN at an early timepoint (5 dpi) with their number peaking between 7-14 dpi, while IgG^+^ PCs emerged at 7 dpi, gradually increasing and peaking at 28 dpi (Figure 1B and S1D). Interestingly, prior to 7 dpi, the ratio of IgA^+^ PCs to IgA^+^ GC B cells was initially much higher than the ratio of IgG^+^ PCs to IgG^+^ GC B cells (Figure 1C), indicating a greater propensity of IgA^+^ B cells to differentiate into PCs than IgG^+^ cells in the earliest days of the immune response. The same was observed in the MLN where, consistent with our previous observations, the early (7 dpi) IgA^+^ PC response was particularly robust in the MLN^24^. Thus, RV infection induces IgA^+^ and IgG^+^ GC responses in the induction sites of the small intestine, with IgA class switched PC arising earlier and more prominently than IgG class switched PC.

**Figure 1.**
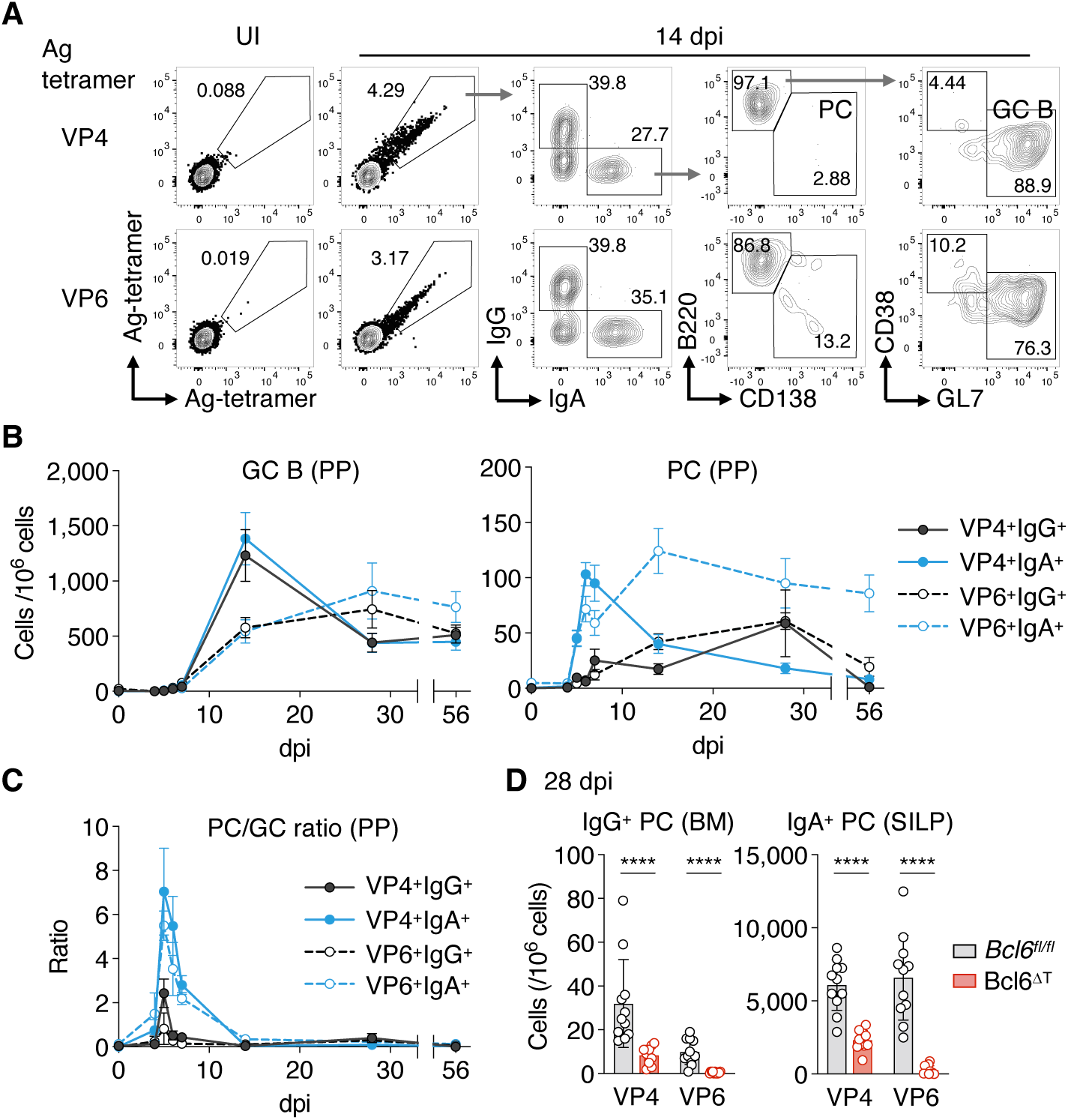
T_FH_ cells are required for IgG and IgA responses to RV. (**A-C**) Flow cytometric analysis of GC B cells and PCs in the PPs after RV infection. Representative flow cytometry profiles showing the gating strategy of RV-specific GC B cells and PCs (A), graphs showing the number of GC B cells and PCs (B) and the ratio of PCs to GC B cells (**C**). Data are pooled from at least 2 independent experiments and shown as mean ± SEM (n = 4- 21). All Abs and Ag-tetramers (VP4 and VP6) in (A) were used for cell surface staining. UI, uninfected. Gating strategy is shown in Figure S1C. (**D**) Absolute number of VP4- and VP6-specific IgG^+^ and IgA^+^ PCs in the BM (bone marrow) and SILP (small intestinal lamina propria) of *Bcl6^fl/fl^* and Bcl6^ΔT^ mice at 28 dpi analyzed by ELISPOT assay. Data are pooled from 3 independent experiments and shown as mean ± SD; each symbol represents one mouse (n = 11); Mann-Whitney test.

We next investigated whether the IgA response to RV requires CD40 ligand (CD40L)- mediated T-cell-help which is critical for a GC reaction and is mainly provided by T_FH_ cells^12^. Administration of an anti-CD40L blocking antibody during the first 2 weeks of RV-infection resulted in a significant reduction in the number of both IgG^+^ and IgA^+^ RV-specific PCs at 28 dpi in the BM and SILP respectively, indicating that both IgG and IgA responses to RV at this timepoint depend on CD40-CD40L interactions (Figure S1F). To investigate the requirement for T_FH_ cells in generating RV-specific IgG^+^ and IgA^+^ PCs, we crossed *Bcl6^fl/fl^* mice and *Cd4^Cre^* mice to create T-cell-specific Bcl6 deficient mice which lack T_FH_ cells (Bcl6^ΔT^ mice - Figure S1G)^25^, and infected them with RV. Compared to littermate controls, the number of VP4- and VP6-specific IgG^+^ and IgA^+^ GC B cells and PCs were significantly decreased in the PPs of Bcl6^ΔT^ mice at 14 dpi (Figure S1H and S1I). Similar results were observed in the MLN (Figure S1J), thus we focused our analysis of induction sites on PPs for the remainder of the study. In terms of effector sites, at 28 dpi, Bcl6^ΔT^ mice had fewer VP4- and VP6-specific IgG^+^ and IgA^+^ PCs than littermate control mice within the PC population in the BM and the SILP respectively (Figure 1D), indicating that establishment of RV-specific IgG^+^ and IgA^+^ PCs at these sites is dependent on T_FH_ cells. Taken together, RV infection induces a precocious IgA^+^ PC response compared to the generation of IgG^+^ PC within the PPs and MLN, and the establishment of both IgG^+^ and IgA^+^ PCs at their effector sites (BM, SILP) is dependent on CD40 signaling and T_FH_.

### RV infection induces gut IgA^+^ LLPCs that produce protective IgA

We next sought to characterize RV-specific IgA^+^ LLPCs. Using recombinant VP4 and VP6 proteins in an enzyme-linked immunosorbent spot (ELISPOT) assay, we detected VP4- and VP6-specific IgA^+^ PCs in the SILP of RV-infected mice but not in naïve mice (Figure 2A and 2B). The number of VP4- and VP6-specific IgA^+^ PCs peaked at 14-28 dpi and was robustly maintained until 200 dpi (Figure 2B). VP4- and VP6-specific IgG^+^ PCs were also detected in the BM but not in the SILP at 7-200 dpi (Figure S2A-C). Therefore, infection with RV ECw strain results in the generation of a durable pool of VP4- and VP6-specific IgA^+^ LLPC in the SILP and IgG^+^ LLPC in the BM.

**Figure 2.**
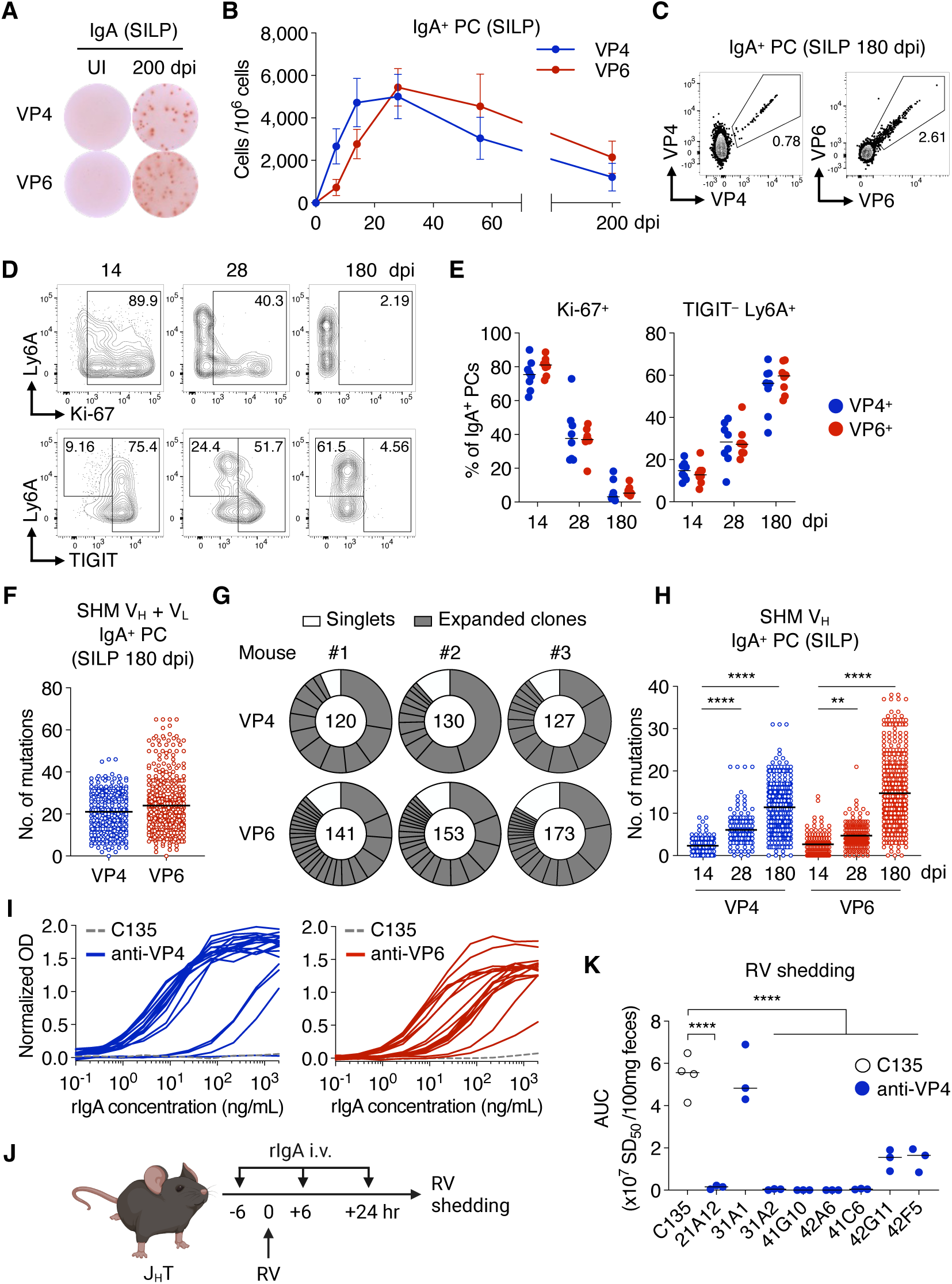
RV infection induces IgA^+^ LLPCs in the gut that protect against infection. (**A and B**) ELISPOT assay detecting VP4- and VP6-specific IgA^+^ PCs after RV infection. Representative ELISPOT image (A) and the number of IgA^+^ PCs in the SILP (B). Data are pooled from at least 2 independent experiments and shown as mean ± SD (n = 8). UI, uninfected. (**C-E**) Flow cytometric analysis of RV-specific IgA^+^ PCs in the SILP after RV infection. Representative plots showing the percentage of VP4^+^ and VP6^+^ among IgA^+^ PCs (C), the FACS profile of Ki-67, Ly6A and TIGIT expression in VP4^+^IgA^+^ PCs (D), and the frequency of Ki-67^+^ and TIGIT^-^Ly6A^+^ cells within the IgA^+^ PC pool (E). Data are pooled from 2 independent experiments; each symbol represents one mouse (n = 8); lines represent the means. Gating strategy is shown in Figure S2D. (**F and G**) Ig repertoire analysis of IgA^+^ PCs in the SILP at 180 dpi with RV. Graph showing the number of SHM (nucleotides, V_H_ + V_L_) and data are pooled from 3 independent experiments; each circle represents one antibody gene, n = 377 (VP4) and n = 467 (VP6) with the number in the middle of the pie chart indicating the number of clones sequenced (F). Pie charts showing distribution of singlets and expanded clones (G). Gating strategy for sorting is shown in Figure S2E. (**H**) The number of SHM (nucleotides, V_H_) in Abs obtained from IgA^+^ PCs in the SILP at 14, 28 and 180 dpi with RV. Data are pooled from 3 independent experiments; each circle represents one antibody gene; VP4: n = 141, 153, 472 (14, 28, 180 dpi, respectively), VP6: n = 131, 160, 526 (14, 28, 180 dpi, respectively). Kruskal-Wallis test with Dunn’s multiple comparison test. Gating strategy for sorting is shown in Figure S2E. (**I**) Ag-binding capacity of recombinant IgA (rIgA) measured by ELISA. rIgAs were cloned from VP4^+^ and VP6^+^IgA^+^ PCs at 180 dpi. C135, anti-SARS-CoV-2 Spike rAb.19 clones of anti-VP4 and 17 clones of anti-VP6 were tested. (**J**) Schematic representation of rIgA administration protocol for (K). J_H_T mice were administrated with 0.5 mg of rIgA at -6, +6 and +24 hours of RV infection. (**K**) Fecal RV shedding analyzed by ELISA and presented as area under the curve (AUC) of SD_50_ from 0 to 7 dpi. Each symbol represents one mouse (n = 3-4); lines represent the means; one-way ANOVA with Holm Sidak’s multiple comparison test.

To characterize the phenotype of gut IgA^+^ PCs, we included VP4 and VP6 tetramers in our flow cytometry panel. Since IgA^+^ PCs retain BCR expression^26^, we used tetramers and anti-IgA Abs for cell surface staining. Employing this approach, we detected VP4^+^ and VP6^+^ IgA^+^ PCs in the SILP of RV-infected mice (Figure 2C and S2D). LLPCs are non-dividing cells (Ki-67^-^) that are derived from a TIGIT^+^ population of PC but ultimately lose expression of TIGIT and adopt the expression of Ly6A^27,28^. Examining these markers, we observed that at 14 dpi, SILP-resident VP4^+^ and VP6^+^ IgA^+^ PCs were largely Ki-67^+^TIGIT^+^Ly6A^-^ but over time (28-180 dpi) they were largely Ki-67^-^TIGIT^-^Ly6A^+^ (Figure 2D and 2E). These data suggest that RV-specific IgA^+^ PCs in the SILP at 14 dpi are proliferative immature PCs (including plasmablasts) and that non-proliferative PCs emerge as early as 28 dpi, becoming the dominant RV-specific PC population over time.

Next, we used *Prdm1*-YFP transgenic mice which report the expression of Blimp1, a transcription factor expressed by PCs, to investigate the clonal structure of the RV-specific IgA^+^ LLPC pool at 180 dpi (Figure S2E). Repertoire analysis of YFP^+^VP4^+^ and YFP^+^VP6^+^ IgA^+^ PCs revealed an accumulation of SHM in both V_H_ and V_L_ genes (Figure 2F) with evidence of clonal expansion (Figure 2G). A time course analysis revealed that the number of SHM in V_H_ genes gradually increased over time (Figure 2H). These data suggest that B cells that have clonally expanded and acquired SHM are preferentially retained in the SILP as IgA^+^ LLPC in response to RV infection. Of note, the number of SHM at 180 dpi was diverse, with a subset of clones possessing a low number of SHMs (14% of VP4^+^ and 8% of VP6^+^ clones had 5 or less SHM in V_H_, Figure 2H). This indicates that at least some PCs generated early in the immune response also develop into LLPCs.

*In vivo* functionality of Abs produced by LLPCs generated following RV infection has never been tested. To examine this, we cloned respective IgH/L from individual VP4^+^ and VP6^+^ IgA^+^ PC at 180 dpi into expression vectors and co-expressed these IgH/L genes with the immunoglobulin J chain (IgJ) in Expi293F cells to produce recombinant dimeric IgAs (rIgAs) (Figure S2F). We validated rIgAs for Ag-binding by enzyme linked immunosorbent assay (ELISA) (Figure 2I) and selected eight anti-VP4 clones based on their binding capacity to Ag for *in vivo* testing. We next administered these anti-VP4 rIgA to B cell deficient J_H_T mice prophylactically, comparing their efficacy in preventing RV infection with a negative control anti-SARS-CoV2 Spike rIgA (clone C135 - Figure 2J). Administered rIgA was successfully secreted into the feces of J_H_T mice (Figure S2G). Seven out of 8 clones reduced RV shedding, and notably 5 clones completely blocked RV shedding, indicating they provided protective immunity against RV infection (Figure 2K). These results demonstrate that RV infection induces clonally expanded populations of IgA^+^ LLPCs in the SILP that produce highly mutated IgA with the capacity to protect the host from infection.

### The development of RV-specific IgA^+^ LLPCs does not require the expression of MHCII on cDCs

Differentiation of T_FH_ cells capable of supporting an IgG^+^ B cell response in the GC is a multistage process that requires sequential Ag presentation by cDCs and B cells^9–11^. However, it is unclear whether the same rules apply to T_FH_ cells that support IgA^+^ B cell responses in induction sites (PPs and MLN) that drain the small intestine. To investigate this, we generated B cell-specific MHCII deficient (MHCII^ΔB^) mice by crossing *H2-Ab1^fl/fl^* mice and *Mb1^Cre^* mice (Figure S3A), and infected these with RV. Consistent with what we observed in Bcl6^ΔT^ mice and with anti-CD40L treatment, compared to littermate controls, at 14 dpi we observed a significant reduction in the numbers of IgG^+^ and IgA^+^ VP4^+^ and VP6^+^ GC B cells in the PPs, and at 28 dpi, we observed a reduction in IgG^+^ and IgA^+^ PCs in the BM and SILP respectively (Figure 3A-C). These data indicate that B cell Ag-presentation is necessary for the development of both IgG^+^ and IgA^+^ RV-specific PCs.

**Figure 3.**
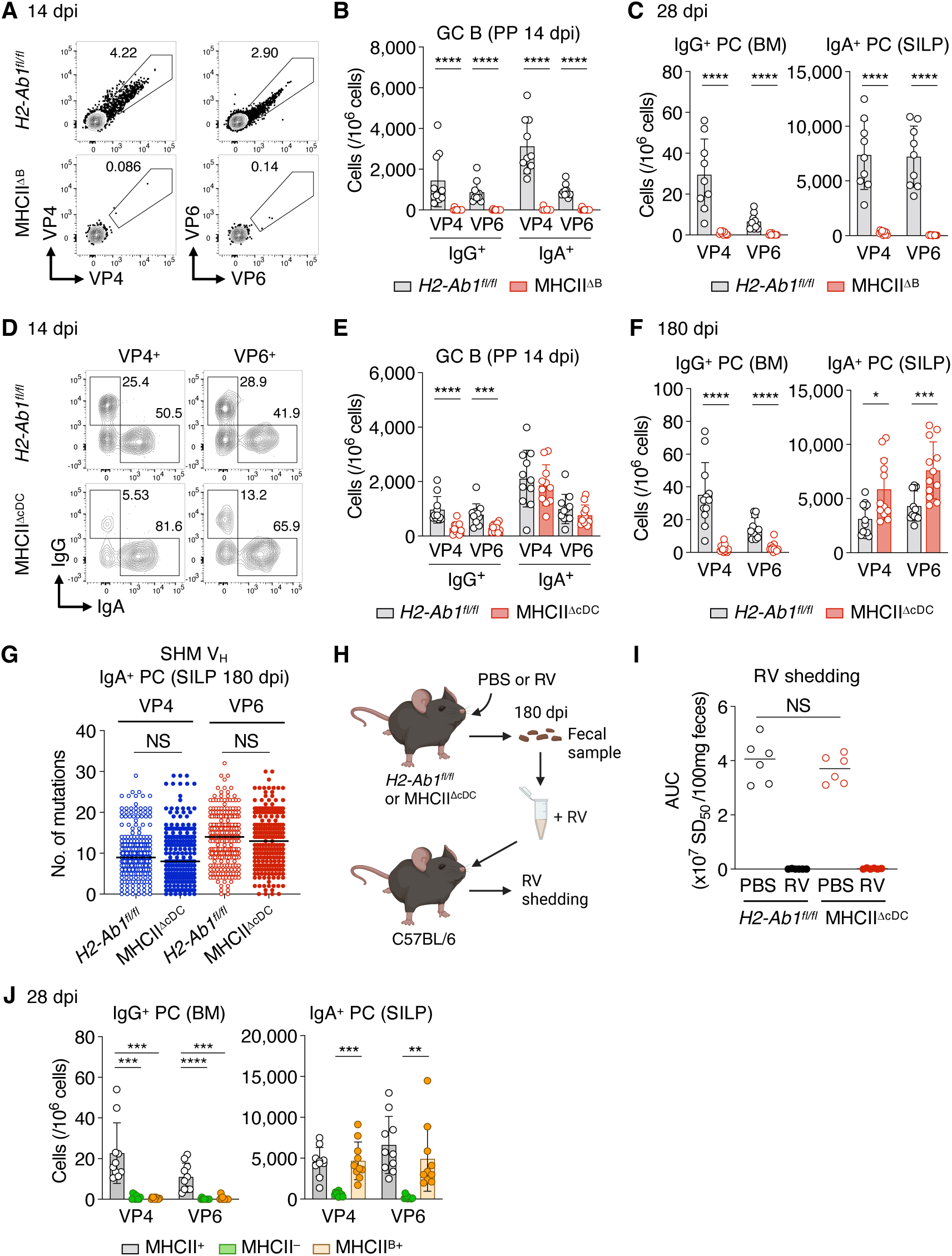
Ag-presentation by B cells but not cDCs is necessary and sufficient for the development RV-specific IgA^+^ LLPCs. (**A and B**) Flow cytometric analysis of the PPs of *H2-Ab1^fl/fl^* and MHCII^ΔB^ mice at 14 dpi with RV. Representative plots showing the percentage of VP4^+^ and VP6^+^ cells among B cells and PCs (A) and graph showing the number of GC B cells (B). Data are pooled from 2 independent experiments and shown as mean ± SD; each symbol represents one mouse (n = 10); Mann-Whitney test. Gating strategy is shown in Figure 1A. (**C**) The number of VP4- and VP6-specific IgG^+^ and IgA^+^ PCs of *H2-Ab1^fl/fl^* and MHCII^ΔB^ mice at 28 dpi analyzed by ELISPOT. Data are pooled from 2 independent experiments and shown as mean ± SD; each symbol represents one mouse (n = 9); Mann-Whitney test. (**D and E**) Flow cytometry analysis of the PPs of *H2-Ab1^fl/fl^*and MHCII^ΔcDC^ mice at 14 dpi with RV. Representative plots showing the percentage of IgG^+^ and IgA^+^ cells among VP4^+^ and VP6^+^ cells (D) and the number of GC B cells (E). Data are pooled from 2 independent experiments and shown as mean ± SD; each symbol represents one mouse (n = 11); Mann-Whitney test. Gating strategy is shown in Figure 1A. (**F**) The number of VP4- and VP6-specific IgG^+^ and IgA^+^ PCs of *H2-Ab1^fl/fl^* and MHCII^ΔcDC^ mice at 180 dpi with RV analyzed by ELISPOT assay. Data are pooled from 2 independent experiments and shown as mean ± SD; each symbol represents one mouse (n = 12); Mann-Whitney test. (**G**) The number of SHM (nucleotides, V_H_) in Abs obtained from VP4^+^ and VP6^+^ IgA^+^ PCs at 180 dpi with RV. Data are pooled from 3 independent experiments; each circle represents one antibody gene, n = 245 (*H2-Ab1^fl/fl^*, VP4), 248 (MHCII^ΔB^, VP4), 245 (*H2-Ab1^fl/fl^*, VP6) and 251 (MHCII^ΔB^,VP6); Mann-Whitney test. (**H and I**) RV neutralizing capacity of secreted Ab in *H2-Ab1^fl/fl^* and MHCII^ΔcDC^ mice. Schematic representation of RV neutralization assay (H) and fecal RV shedding analyzed by ELISA and presented as AUC of SD_50_ from 0 to 7 dpi (I). Each symbol represents one mouse (n = 6); lines represent mean. Kruskal-Wallis test with Dunn’s multiple comparison test. (**J**) The number of VP4- and VP6-specific IgG^+^ and IgA^+^ PCs of MHCII^+^ (MHCII sufficient), MHCII^-^ (MHCII deficient) and MHCII^B+^ (MHCII only in B cells) mice at 28 dpi with RV analyzed by ELISPOT assay. CD4^+^ T cells purified from uninfected C57BL/6 mice were transferred to MHCII^-^ and MHCII^B+^ mice 1 day before the infection to reconstitute the CD4^+^ T cell pool. Data are pooled from 2 independent experiments and shown as mean ± SD; each symbol represents one mouse (n = 10); Kruskal-Wallis test with Dunn’s multiple comparison test.

To assess whether cDC-mediated Ag-presentation is required for the generation of either IgG^+^ or IgA^+^ RV-specific PCs, we next generated cDC-specific MHCII deficient (MHCII^ΔcDC^) mice by crossing *H2-Ab1^fl/fl^* mice and *Zbtb46^Cre^* mice (Figure S3B), analyzing the B cell response after RV infection. Compared to littermate controls, MHCII^ΔcDC^ mice exhibited a reduced number of VP4^+^ and VP6^+^ IgG^+^ GC B cells in the PPs at 14 dpi and IgG^+^ PCs in the BM at 28 and 180 dpi (Figure 3D-F and S3C). However, unexpectedly, MHCII^ΔcDC^ mice did not exhibit a reduced number of VP4^+^ and VP6^+^ IgA^+^ GC B cells in the PPs nor VP4^+^ and VP6^+^ IgA^+^ PCs in the SILP (Figure 3D-F and S3C). Moreover, repertoire analysis revealed that the number of V_H_ SHM of VP4^+^ and VP6^+^ IgA^+^ PCs at 180 dpi was comparable between MHCII^ΔcDC^ mice and littermate controls (Figure 3G). To assess the neutralizing capacity of secreted IgA Abs generated in the absence of MHCII on cDC, we collected fecal samples from MHCII^ΔcDC^ mice vs littermate controls either uninfected or RV-infected (180 dpi), incubated these fecal samples with live RV and used this mixture to infect naïve C57BL/6 (B6) mice (Figure 3H). Compared to the fecal samples from naïve mice, those from MHCII^ΔcDC^ RV-infected mice completely prevented viral shedding into the feces at a level comparable to RV-infected littermate control mice (Figure 3I), indicating that secreted Abs generated in MHCII^ΔcDC^ mice have the capacity to neutralize RV. These data demonstrate that, unlike the systemic IgG response to RV, the development of functional RV- specific IgA^+^ LLPCs does not require the expression of MHCII on cDCs.

Given that we had observed that the expression of MHCII on B cells was required for the generation of RV-specific IgA^+^ LLPCs in the SILP, yet MHCII expression on cDCs was dispensable, we next asked whether expression of MHCII on B cells is sufficient to support the IgA^+^ LLPC response to RV infection. To test this, we generated mice that allow conditional expression of MHCII specifically and only in B cells by crossing *H2-Ab1^loxP-Stop-loxP^* (*H2-Ab1^LSL^*) mice^29^ with *Cd19^Cre^* mice. We compared the RV-specific IgG and IgA responses in *H2-Ab1^LSL/LSL^Cd19^Cre^* (MHCII^B+^) mice with MHCII-sufficient *H2-Ab1^LSL/+^Cd19^Cre^* (MHCII^+^) mice – a positive control, and MHCII-deficient *H2-Ab1^LSL/LSL^* (MHCII^-^) mice – a negative control (Figure S3D). Since T cells are absent in MHCII^-^ and MHCII^B+^ mice due to the lack of the thymic selection^29^, we supplemented MHCII^-^ and MHCII^B+^ mice with an adoptive transfer of polyclonal CD4^+^ T cells purified from uninfected B6 mice. As expected, at 28 dpi MHCII^-^ mice exhibited a significantly reduced number of VP4^+^ and VP6^+^ IgG^+^ and IgA^+^ PCs in the BM and SILP respectively compared to MHCII^+^ mice. Remarkably, while MHCII^B+^ mice were unable to support an IgG^+^ PC response, these mice harbored a similar number of VP4- and VP6-specific IgA^+^ PCs in the SILP compared to MHCII^+^ mice (Figure 3J). Thus, we conclude that B cell intrinsic MHCII expression is necessary and sufficient for the accumulation of VP4- and VP6-specific IgA^+^ PCs in the SILP following RV infection.

### Characterization of RV-specific CD4^+^ T cells

It has been proposed that specialized subsets of T_FH_ cells have different roles in supporting isotype specific Ab responses^11^ and that the phenotypic fate of CD4^+^ T cells can be imprinted during Ag presentation^30^. Our finding of different APC requirements for the generation of VP4- and VP6- specific IgG^+^ and IgA^+^ LLPCs prompted us to characterize RV-induced VP4- and VP6-specific T_FH_ cells. To investigate this, it was necessary to distinguish RV-specific T_FH_ cells from microbiota and food Ag-specific T_FH_ cells. We therefore mapped the antigenic peptides of RV ECw strain VP4 and VP6 proteins that provoke a T_FH_ cell response by using an activation induced marker (AIM) assay as a readout^31^ in response to an overlapping VP4 and VP6 peptide library (Figure S4A and S4B). Peptide pools that had >1% AIM positivity were deconvoluted and individual peptides from these pools were further tested for AIM positivity (Figure S4C and S4D). This screening process identified four VP4 antigenic peptides (p51, p52, p110 and p120) and three VP6 antigenic peptides (p58, p61 and p62) for T_FH_ cells. Next, we generated fluorochrome-labeled peptide:I-A^b^ tetramers presenting the identified VP4 and VP6 peptides and tested them for staining PP- and MLN-derived CD4^+^ T cells of RV-infected B6 mice at 7 dpi. After a sensitive magnetic sort-based cell enrichment method^32^, we identified VP4-p110:I-A^b^ and VP6-p61:I-A^b^ tetramers as suitable reagents for detecting RV-specific T_FH_ cells (Figure S4E and S4F). Hereafter, we used these two tetramers for subsequent analyses, referring to them as VP4:I-A^b^ and VP6:I-A^b^.

Using these tetramers, we characterized VP4- and VP6-specific T_FH_ cell subsets in RV infected mice at 7 dpi. We purified CD4^+^ T cells bound by VP4:I-A^b^ and VP6:I-A^b^ tetramers (Figure 4A and S4G) as well as total CD4^+^ T cells from pooled PPs and MLNs of RV infected B6 mice and subjected the admixture to single cell RNA-sequencing (scRNA-seq) analysis. We identified 11 distinct clusters of CD4^+^ T cells, each annotated based on selected marker genes. Visualizing these clusters by uniform manifold approximation and projection (UMAP), we observed that VP4:I-A^b+^ and VP6:I-A^b+^ cells were dominated by T_H_1 cells defined by expression of *Tbx21* (encodes T-bet), *Ifng*, *Cxcr3*, *Cxcr6 and Selplg* (encodes PSGL-1) and T_FH_ cells defined by expression of *Bcl6* and *Cxcr5* (Figure 4B, 4C and S4H). Re-clustering of VP4:I-A^b+^ and VP6:I- A^b+^ cells alone further revealed 7 clusters based on selected marker genes: C0 (naïve), C1-4 (T_H_1) and C5-6 (T_FH_) (Figure 4D, 4E and S4I). Differential gene expression analysis of C5 vs C6 T_FH_ cells identified up- and down-regulated genes (Figure S4J); gene set enrichment analysis (GSEA) revealed that C6 T_FH_ cells express significantly higher levels of genes related to c-Myc, mTORC1 and IL-2–STAT5 signaling compared to C5 T_FH_ cells (Figure S4K). These pathways have been linked to T_H_1 cell differentiation, partly due to the promotion of T-bet expression, a master regulator of T_H_1 cells^33,34^. Consist with these observations, we found that C6 T_FH_ cells uniquely expressed T_H_1 genes, *Tbx21*, *Ifng* and *Cxcr3*; however, they did not express *Cxcr6* which is highly expressed by all T_H_1 clusters (Figure 4E). In summary, our scRNA-seq analysis identified a subset of RV-specific T_FH_ cells exhibiting type 1 characteristics. Given their dual expression *of Tbx21* and *Bcl6*, we refer to C6 cells as T_FH_1 cells.

**Figure 4.**
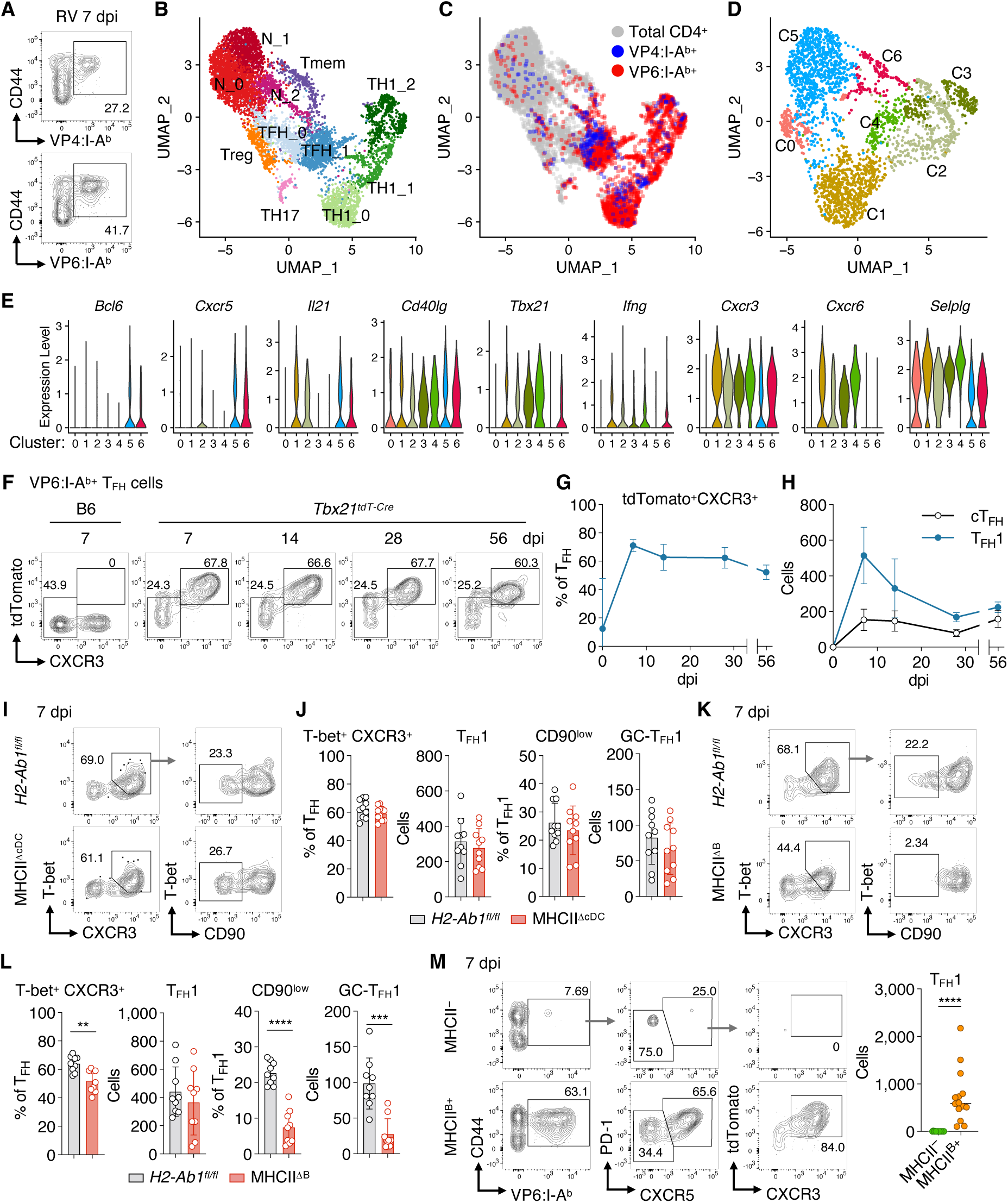
scRNA-seq identifies RV-specific T_FH_1 cells, the development of which requires unconventional priming by B cells. (**A**) Representative flow cytometry plots showing the binding of VP4:I-A^b^ and VP6:I-A^b^ tetramers in CD45^+^Dump^-^CD3^+^CD4^+^ T cells from C57BL/6 mice at 7 dpi with RV. Peptide:I-A^b+^ cells were enriched from pooled PP and MLN cells by magnetic sort and used for analysis. Gating strategy is shown in Figure S4G. (**B-D**) scRNA-seq analysis of total and RV-specific CD4^+^ T cells in PPs and MLNs of C57BL/6 mice at 7 dpi with RV. UMAP plots of total, VP4:I-A^b+^ and VP6:I-A^b+^ CD4^+^ T cells (4417, 372 and 2529 cells, respectively) (B), distribution of total, VP4:I-A^b+^ and VP6:I-A^b+^ CD4^+^ T cells on UMAP (C) and UMAP plots of VP4:I-A^b+^ and VP6:I-A^b+^ CD4^+^ T cells only (D). Data are pooled from 2 independent experiments. (**E**) Violin plots showing expression levels of T_FH_ and T_H_1 genes across all clusters in (D). (**F-H**) Flow cytometric analysis of T_FH_ cells in *Tbx21^tdT-Cre^* mice following RV infection. Representative plots showing the frequencies of tdTomato^+^CXCR3^+^ T_FH_1 and tdTomato^-^CXCR3^-^ cT_FH_ among VP6:I-A^b+^ T_FH_ cells at the indicated time points (F), frequency of tdTomato^+^CXCR3^+^ T_FH_1 among VP6:I-A^b+^ T_FH_ cells (G) and the absolute number of VP6:I-A^b+^ T_FH_1 and VP6:I-A^b+^ cT_FH_ cells (H). Cells were processed and gated as in (A). Data are pooled from 2 independent experiments and shown as mean ± SD (n = 6-8). (**I and J**) Flow cytometric analysis of T_FH_1 cells in *H2-Ab1^fl/fl^* and MHCII^ΔcDC^ mice at 7 dpi. Representative plots showing the percentage of T-bet^+^CXCR3^+^ T_FH_1 cells among T_FH_ cells and the percentage of CD90^low^ cells among T_FH_1 cells (I) and graphs showing the frequencies and numbers of T-bet^+^CXCR3^+^ T_FH_1 cells and CD90^low^ GC-T_FH_1 cells (J). Cells were processed and gated as in (A). Data are pooled from 2 independent experiments and shown as mean ± SD; each symbol represents one mouse (n = 10). (**K and L**) Flow cytometric analysis of T_FH_1 cells in *H2-Ab1^fl/fl^* and MHCII^ΔΒ^ mice at 7 dpi with RV, presented as in (I and J). Data are pooled from 2 independent experiments and shown as mean ± SD; each symbol represents one mouse (n = 9); Mann-Whitney test. (**M**) Flow cytometric analysis of T_FH_1 cells in MHCII^-^ and MHCII^B+^ mice at 7 dpi with RV. Total CD4^+^ T cells purified from uninfected *Tbx21^tdT-Cre^* mice were transferred 1 day before the infection to reconstitute the CD4^+^ T cell pool. Representative plots showing the gating for tdTomato^+^CXCR3^+^ T_FH_1 cells and graph showing the number of T_FH_1 cells. Data are pooled from 3 independent experiments; each symbol represents one mouse (n = 14); lines represent mean; Mann-Whitney test. Cells were processed as in (A).

To further investigate RV-induced T_FH_1 cells, we utilized *Tbx21^tdTomato-T2A-Cre^* knock-in mice (*Tbx21^tdT-Cre^*), whereby tdTomato (tdT) faithfully reports endogenous *Tbx21* expression^35^. Following magnetic enrichment of VP6:I-A^b^ tetramer binding cells (the most dominant CD4^+^ T cell binder in RV infected mice - Figure S4F) from pooled PP and MLN, we observed both VP6:I- A^b+^CXCR3^-^CXCR5^+^PD-1^+^ conventional T_FH_ (cT_FH_) cells that do not express tdTomato as well as VP6:I-A^b+^CXCR3^+^CXCR5^+^PD-1^+^ T_FH_1 cells that express tdTomato, with both cT_FH_ and T_FH_1 peaking in the PPs and MLN at 7 dpi and maintained until at least 56 dpi (Figure 4F-H). Furthermore, high parameter flow cytometry analysis using *Tbx21^tdT-Cre^* mice validated that, unlike T_H_1 cells, T_FH_1 cells express BCL6 and exhibit lower expression of CXCR6 and PSGL-1 compared to T_H_1 cells (Figure S4L). We conclude that RV infection induces distinct cT_FH_ and T_FH_1 cells in the PPs and MLN as early as 7 dpi.

### The development of RV-specific T_FH_1 cells does not require the expression of MHCII on cDCs

We observed that the expression of MHCII on B cells was sufficient for the generation of RV-specific IgA^+^ LLPCs (Figure 3J). If the generation of RV-specific IgA^+^ LLPCs is supported by T_FH_1 cells, we hypothesize that these T_FH_1 cells would likewise be generated independent of MHCII expression by cDCs. To test this, we examined RV-specific T_FH_1 cell development in MHCII^ΔcDC^ mice. Although the sensitivity was lower than the *Tbx21^tdT-Cre^* reporter, we successfully detected T_FH_1 cells among the VP6:I-A^b+^ RV-specific T_FH_ population in pooled PPs and MLN by intracellular staining of T-bet in combination with CXCR3 staining (Figure 4I). We also incorporated staining for CD90 since low expression of this glycoprotein has been reported as a marker of GC-resident T_FH_ (GC-T_FH_)^36^. We observed a similar proportion and number of both T-bet^+^CXCR3^+^CD90^high^ and T-bet^+^CXCR3^+^CD90^low^ T_FH_1 cells in MHCII^ΔcDC^ mice compared to littermate controls (Figure 4I and 4J), indicating that Ag-presentation by cDCs is dispensable for T_FH_1 cell development, including GC-resident T_FH_1. The same analysis in MHCII^ΔB^ mice identified that the proportion of T-bet^+^ CXCR3^+^ T_FH_1 cells was decreased in MHCII^ΔB^ mice, although their absolute number was not significantly changed (Figure 4K and 4L). However, both the proportion and the number of CD90^low^ T_FH_1 cells was significantly decreased in MHCII^ΔB^ mice (Figure 4K and 4L). These data suggest that Ag-presentation by B cells, but not cDC, is necessary for induction and maturation of GC-resident T_FH_1 cells.

To investigate the sufficiency of B cell Ag presentation for T_FH_1 cell development, we performed the same experiment as described in Figure 3J whereby we used the conditional MHCII expression system. Following transfer of polyclonal CD4 T cells purified from uninfected *Tbx21^tdT-Cre^* mice to MHCII^-^ and MHCII^B+^ mice, we subsequently infected these mice with RV, enumerating the number of VP4- and VP6-specific donor-derived T_FH_1 cells. At 7 dpi, we detected tdTomato^+^CXCR3^+^ T_FH_1 cells in pooled PP and MLN of MHCII^B+^ recipients but not in MHCII^-^ recipients (Figure 4M). Therefore, like RV-specific IgA^+^ LLPCs, the expression of MHCII by B cells is both necessary and sufficient for the accumulation of VP6-specific T_FH_1 cells in the induction sites (PPs and MLN) in response to RV infection.

### The development of RV-specific IgA^+^ LLPCs requires IFNγ production by T_FH_1 cells

Although previous studies have shown a requirement for T-bet-expressing T_FH_1 cells in different infection models^37–39^, the role for T_FH_1 in supporting IgA responses is not known. Since neither RV-specific T_FH_1 nor IgA^+^ LLPCs rely on MHCII expression by cDCs for their respective accumulation in induction sites and the effector site, we hypothesized that T_FH_1 cells may be uniquely required for generating an IgA response to RV. To test this hypothesis, we generated T_FH_1 deficient mice. Specifically, we made mixed-BM chimeras, reconstituting lethally irradiated *Tcrb^-/–^* mice with an 80:20 mixture of *Tcrb^-/–^* BM (80%) with either 20% *Bcl6^+/+^Tbx21^tdT-Cre^* BM or 20% *Bcl6^fl/fl^Tbx21^tdT-Cre^* BM (Figure 5A). The resulting chimeric mice will harbor T-bet^+^ αβ T cells that can either express Bcl6 (*Bcl6^+/+^Tbx21^tdT-Cre^*, control chimeras) or cannot express Bcl6 (*Bcl6^fl/fl^Tbx21^tdT-Cre^*, experimental chimeras) ((Figure S5A and (S5B). After RV infection, control and experimental chimeras exhibited comparable numbers of IgG^+^ and IgA^+^ GC B cells and PCs in the PPs at 14 dpi and of IgG^+^ PCs in the BM at 70 dpi (Figure 5B-D). However, the experimental chimeras exhibited a significant reduction in the number of VP4- and VP6-specific IgA^+^ PCs in the SILP at 70 dpi (Figure 5D). These data demonstrate that while T_FH_1 cells are not required for the initial generation of an IgA response in the induction site (PPs) they are essential for the accumulation and/or maintenance of IgA^+^ LLPCs in the SILP in response to RV infection.

**Figure 5.**
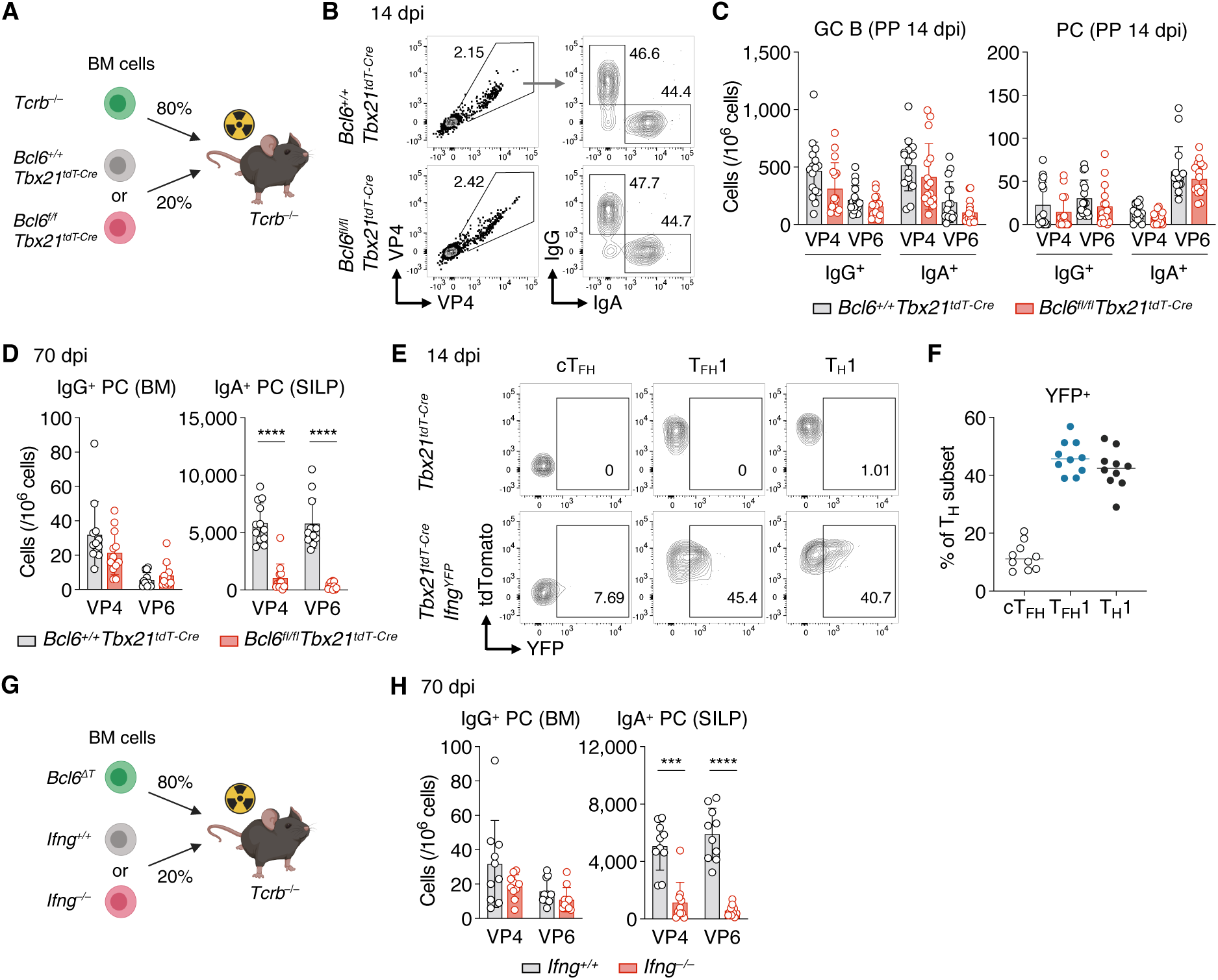
T_FH_1 cells are required for the development of IgA^+^ LLPCs by providing IFNγ. (**A**) Schematic representation of mixed-BM chimeras for (B-D). After BM reconstitution, chimeras were infected with RV. (**B and C**) Flow cytometric analysis of mixed-BM chimeras at 14 dpi. Representative plots showing the percentage of IgG^+^ and IgA^+^ cells among VP4^+^ cells (B) and graphs showing the number of GC B cells and PCs (C). Data are pooled from 3 independent experiments and shown as mean ± SD; each symbol represents one mouse (n = 16). Gating strategy is shown in Figure 1A. (**D**) Enumeration of VP4- and VP6-specific IgG^+^ and IgA^+^ PCs in mixed-BM chimeras at 70 dpi. ELISPOT data are pooled from 2 independent experiments and shown as mean ± SD; each symbol represents one mouse (n = 12); Mann-Whitney test. (**E and F**) Flow cytometric analysis of *Tbx21^tdT-Cre^* and *Tbx21^tdT-Cre^Ifng^YFP^* reporter mice at 14 dpi. Representative plots showing the percentage of YFP^+^ cells among VP6:I-A^b+^ cT_FH_, T_FH_1, and T_H_1 cells (E) and graph showing the frequencies of YFP^+^ cells (F). Data are pooled from 2 independent experiments; each symbol represents one mouse (n = 10); lines represent means. Cells were processed and gated as Figure 4F. T_H_1 cells were defined as tdTomato^+^ non-T_FH_ cells gated as in (Figure S4G. (**G**) Schematic representation of mixed-BM chimeras for (H). After reconstitution, the chimeras were infected with RV. (**H**) Enumeration of VP4- and VP6-specific IgG^+^ and IgA^+^ PCs in mixed-BM chimeras at 70 dpi. ELISPOT data are pooled from 2 independent experiments and shown as mean ± SD; each symbol represents one mouse (n = 10); Mann-Whitney test.

Given that T_FH_1 cells are not required for initiating the GC reaction or PC generation in the PPs, we hypothesized that T_FH_1 cells express B cell stimulating factor(s) that instruct the accumulation of IgA^+^ PCs in the SILP. Our scRNA-seq data identified that *Ifng* is uniquely expressed by T_FH_1 cells compared to cT_FH_ cells (Figure 4E). To assess the role of IFNγ in the RV-specific IgA response, we crossed *Tbx21^tdT-Cre^*mice with *Ifng^YFP^* mice^40^, creating double *Tbx21^tdT-Cre^Ifng^YFP^* reporter mice, and examined YFP expression in different VP6-specific CD4^+^ T cell subsets at 14 dpi. We observed YFP positivity in tdTomato-positive VP6:I-A^b+^ T_FH_1 cells compared to tdTomato-negative VP6:I-A^b+^ cT_FH_ cells, and the frequency of YFP^+^ cells among VP6:I-A^b+^ T_FH_1 cells was similar to the frequency of YFP^+^ cells among VP6:I-A^b+^ T_H_1 cells (Figure 5E and 5F). These findings suggests that T_FH_1 cells are the dominant IFNγ producers among T_FH_ cells in response to RV.

To assess the role of T_FH_-derived IFNγ, we generated mixed-BM chimeras by reconstituting lethally irradiated *Tcrb^-/–^* mice with an 80:20 mixture of *Bcl6^fl/fl^Cd4^Cre^*BM and either *Ifng^+/+^* or *Ifng^-/–^* BM (Figure 5G). At 70 dpi, compared to *Ifng^+/+^* control chimeras, *Ifng^-/–^*experimental chimeras showed a significant reduction in the number of VP4- and VP6-specific IgA^+^ PCs in the SILP while the number of IgG^+^ PCs in the BM was unaffected (Figure 5H). From these data, we conclude that T_FH_1-derived IFNγ is essential for the development of IgA^+^ LLPCs in the SILP but not IgG^+^ LLPC in the BM.

### An IFNγR-CXCR3 axis is necessary for the accumulation of IgA^+^ PCs in the SILP

A role for IFNγ in supporting the development of SILP-resident IgA^+^ PCs has never been described. To investigate the mechanism for how IFNγ controls IgA response to RV, we first assessed whether IFNγ directly affects B cells. To this end, we generated mixed-BM chimeras by reconstituting lethally irradiated B cell deficient J_H_T mice with a 50:50 mixture of CD45.1- *Ifngr1*^+/+^ and CD45.2-*Ifngr1*^-/–^ BM (Figure 6A), allowing us to assess the role of B cell-intrinsic IFNγ-receptor (IFNγR) signaling. After RV infection, we analyzed the chimerism of CD45.1^+^ and CD45.2^+^ target cells (VP4- and VP6-specific GC B cells and PCs) vs naïve B cells. At 14 dpi, we did not observe in impact of cell-intrinsic IFNγR expression on the accumulation of VP4/VP6-specific IgG^+^ or IgA^+^ GC B cells nor IgG^+^ or IgA^+^ PCs in the PPs (Figure 6B and (S6A). However, examining the SILP, we observed a reduction in *Ifngr1*^-/–^ VP4- and VP6-specific IgA^+^ PCs compared to *Ifngr1*^+/+^ VP4- and VP6-specific IgA^+^ PCs at 14 dpi, which was even more profound at 70 dpi, whereas BM-resident IgG^+^ PCs were unaffected by IFNγR deficiency (Figure 6C-F, (S6B-D). These results indicate that while B cell intrinsic IFNγR-signaling is not required within the PP for the generation of IgA^+^ GC B cells or PCs, the accumulation of IgA^+^ LLPCs in the SILP is highly sensitive to cell-intrinsic IFNγR-signaling.

**Figure 6.**
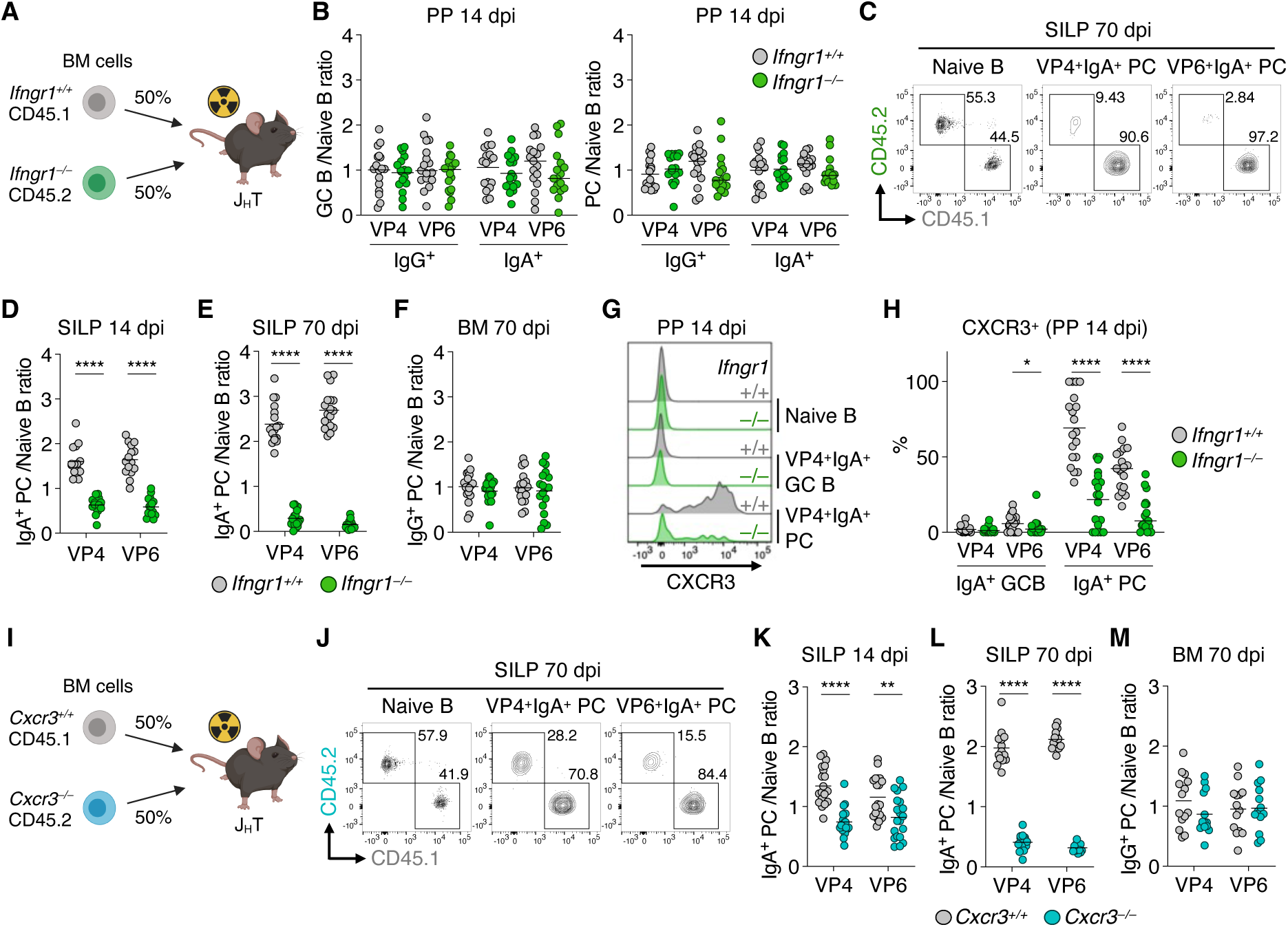
A B cell intrinsic IFN*γ*R-CXCR3 axis is required for the accumulation of IgA^+^ PCs to the SILP. (**A**) Schematic representation of mixed-BM chimeras for (B-H). After reconstitution, chimeras were infected with RV. (**B-F**) Flow cytometric analysis of mixed-BM chimeras. Graphs showing chimerism at 14 dpi in the PPs on a per-mouse basis, expressed as the ratio of GC B cells or PCs to naïve B cells of *Ifngr1*^+/+^ and *Ifngr1*^-/–^ cells (B). Representative plots showing the percentage of CD45.1^+^ (*Ifngr1*^+/+^) and CD45.2^+^ (*Ifngr1*^-/–^) cells among naive B cells and IgA^+^ PCs in the SILP at 70 dpi (C). Graphs showing the chimerism in SILP IgA^+^ PCs at 14 dpi (D), in SILP IgA^+^ PCs at 70 dpi (E) and in BM IgG^+^ PCs at 70 dpi (F). Data are pooled from at least 2 independent experiments; each symbol represents one mouse (n = 17 for B, n = 15 for D, n = 18 for E and F); lines represent mean; gating strategy is shown in Figure 1A, (S6B and (S6C; Wilcoxon test was used for pairwise (intra-chimera) comparisons. (**G and H**) Flow cytometric analysis of mixed-BM chimeras at 14 dpi. Representative histograms showing the expression of CXCR3 in *Ifngr1*^+/+^ and *Ifngr1*^-/–^ cells in the PPs (G) and graph showing the percentage of CXCR3^+^ cells among IgA^+^ GC B cells and PCs in the PPs (H). Data are pooled from 3 independent experiments; each symbol represents one mouse (n = 19); lines represent mean; gating strategy is shown in Figure 1A; Wilcoxon test. (**I**) Schematic representation of mixed-BM chimeras for (J-M). After reconstitution, the chimeras were infected with RV. (**J-M**) Flow cytometry analysis of the mixed-BM chimeras. Representative plots showing the percentage of CD45.1^+^ (*Cxcr3*^+/+^) and CD45.2^+^ (*Cxcr3*^-/–^) cells among naive B cells and IgA^+^ PCs in the SILP at 70 dpi (J) and graphs showing the chimerism of *Cxcr3*^+/+^ and *Cxcr3*^-/–^ cells among SILP IgA^+^ PCs at 14 dpi (K), at 70 dpi (L) and among BM IgG^+^ PCs at 70 dpi (M). Data are pooled from at least 2 independent experiments; each symbol represents one mouse (n = 20 for K, n = 14 for L and M); lines represent means; gating strategy is shown in (Figure S6B and (S6C; Wilcoxon test was used for pairwise (intra-chimera) comparisons.

Based on the above results, we hypothesized that IFNγR-signaling controls the migration of RV-specific IgA^+^ PCs by regulating the expression of homing receptors. Given that IFNγR-signaling is known to upregulate the chemokine receptor CXCR3 in T cells and NK cells^41,42^ and that CXCR3 is required for IgA^+^ PC localization in the lung following IAV infection^43^, we postulated that a potential candidate for homing of VP4/VP6-specific IgA^+^ PCs to the SILP in response to RV infection could be CXCR3. Indeed, we found that CXCR3 was highly expressed on IgA^+^ PCs but not on IgA^+^ GC B cells in the PPs at 14 dpi (Figure 6G). In addition, CXCR3 expression was reduced on *Ifngr1*^-/–^ derived IgA^+^ PCs compared to *Ifngr1*^+/+^ derived IgA^+^ PCs, and the majority of CXCR3^+^IgA^+^ PCs were derived from *Ifngr1*^+/+^ cells in RV-infected 50:50 BM chimeras (Figure 6G and 6H), indicating that B cell intrinsic IFNγ-signaling drives CXCR3 expression on RV-specific IgA^+^ PCs.

To assess the role of CXCR3 on the homing of VP4- and VP6-specific IgA^+^ PCs to the SILP in response to RV infection, we generated mixed-BM chimeras by reconstituting lethally irradiated J_H_T mice with a 50:50 mixture of CD45.1-*Cxcr3*^+/+^ and CD45.2-*Cxcr3^-/–^*BM (Figure 6I), and infected these chimeric mice with RV. We observed a significant reduction in SILP IgA^+^ PCs derived from *Cxcr3^-/–^*cells compared to SILP IgA^+^ PCs derived from *Cxcr3^+/+^*cells in the SILP at 14 and 70 dpi, whereas accumulation of VP4- and VP6-specific IgG^+^ PCs in the BM was unaffected by the expression of *Cxcr3* (Figure 6J-M), phenocopying the 50:50 *Ifngr1*^-/–^ BM chimeras. These data indicate that B cell intrinsic IFNγR and CXCR3 are both required for the localization of RV-specific IgA^+^ PCs to the SILP.

### Differential requirement of cDC-intrinsic MHCII and Tfh1 for the development of influenza-specific IgA^+^ LLPCs

We next determined if our findings with RV infection were generalizable to extraintestinal mucosal viral infections. In a recent study by Iwasaki and colleagues, intranasal infection with IAV in mice was shown to generate lung-resident IgA^+^ PCs which were detected for at least 5 weeks post-infection^43^. However, the role of T_FH_ cells in the IgA response following IAV infection is not known. We therefore intranasally infected mice with a mouse-adapted IAV, A/PR/8/34 H1N1 (PR8), and used recombinant PR8 hemagglutinin (HA) ((Figure S7A) to measure HA-specific PCs by ELISPOT in different tissues. We detected HA-specific IgA^+^ and IgG^+^ PCs in the nasal mucosa (NM), lung and BM in PR8-infected B6 mice at 180 dpi (Figure 7A and 7B), demonstrating that IAV infection induces IgA^+^ LLPCs in both the NM and lung as well as in the BM.

**Figure 7.**
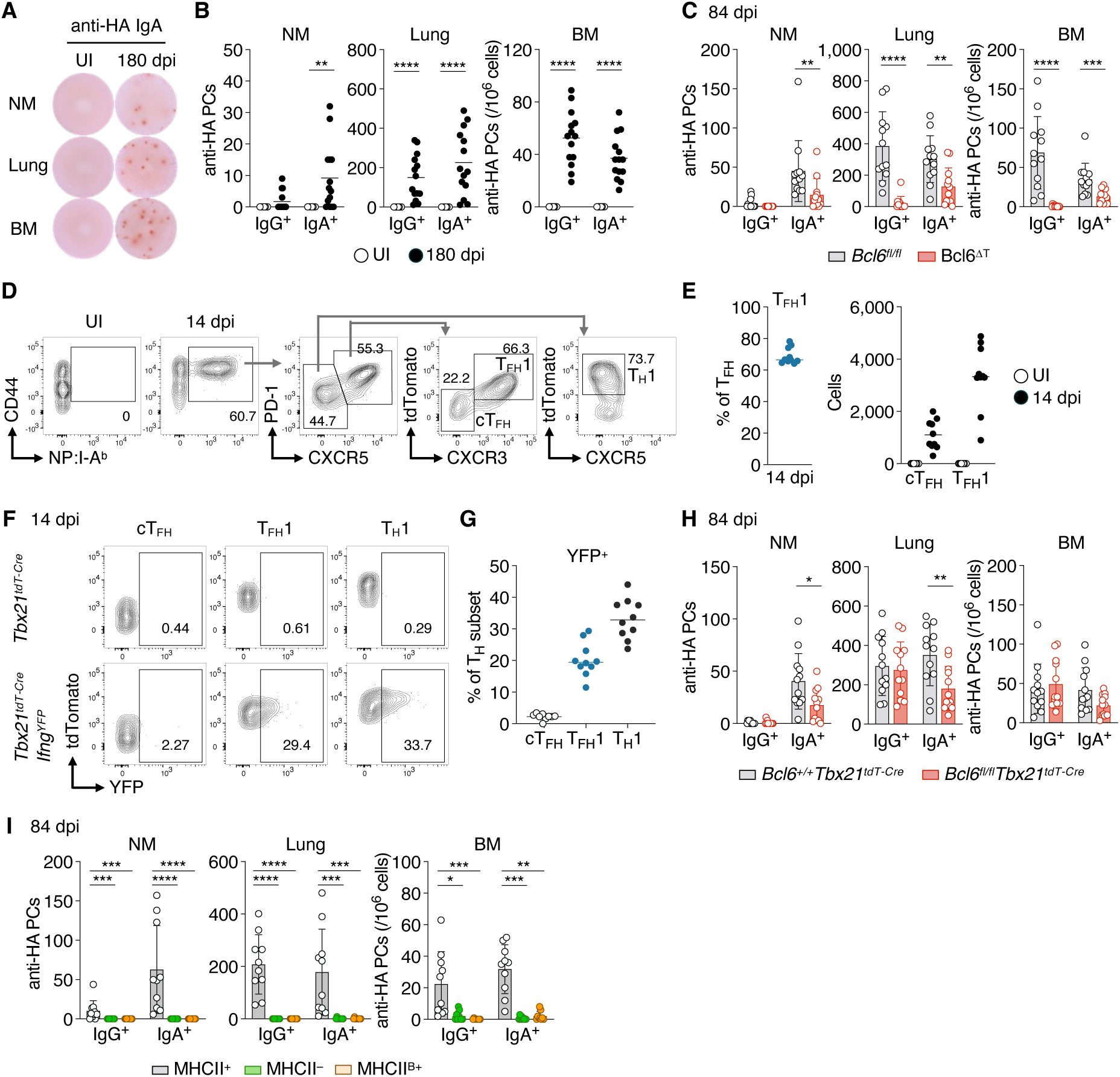
The development of IAV-specific IgA^+^ LLPCs requires T_FH_1 cells. (**A and B**) ELISPOT assay detecting HA-specific PCs in C57BL/6 mice after PR8 infection. Representative ELISPOT image (A) and the number of IgG^+^ and IgA^+^ PCs in the nasal mucosa (NM), lung and BM (B). UI, uninfected. Data are pooled from 2 independent experiments; each symbol represents one mouse (n = 6 for UI, n =8 for 180 dpi); lines represent means; Mann-Whitney test. (**C**) The number of HA-specific IgG^+^ and IgA^+^ PCs at 84 dpi in *Bcl6^fl/fl^* and Bcl6^ΔT^ mice. ELISPOT data are pooled from 2 independent experiments and shown as mean ± SD; each symbol represents one mouse (n = 12); Mann-Whitney test. (**D-G**) Flow cytometric analysis of mediastinal LN cells in *Tbx21^tdT-Cre^* and *Tbx21^tdT-Cre^Ifng^YFP^* mice unimmunized (UI) or at 14 dpi. Representative flow cytometry plots showing the frequencies of tdTomato^+^CXCR3^+^ T_FH_1, tdTomato^-^CXCR3^-^ cT_FH_ and tdTomato^+^ T_H_1 cells among NP:I-A^b+^ population (D), graphs showing the frequencies of tdTomato^+^CXCR3^+^ T_FH_1 cells among NP:I-A^b+^ T_FH_ cells, and the absolute number of NP:I-A^b+^ cT_FH_ and T_FH_1 cells (E). Representative plots showing the percentage of YFP^+^ cT_FH_, T_FH_1 and T_H_1 cells gated as in (D) among NP:I-A^b+^ cells (F), and graph showing the frequencies of YFP^+^ cells (G). Cells were processed and gated as in (Figure S4G using NP_311-325_:I-A^b^ tetramer. Data are pooled from 2 independent experiments and shown as mean ± SD (n = 9 for UI, n = 10 for 14 dpi). (**H**) ELISPOT analysis of 80:20 mixed-BM chimeras employing *Tcrb^-/–^*BM and either *Bcl6^+/+^Tbx21^tdT-Cre^ or Bcl6^fl/fl^Tbx21^tdT-Cre^* BM as shown in Figure 5A. Number of HA-specific IgG^+^ and IgA^+^ PCs in BM chimeras at 84 dpi. Data are pooled from 2 independent experiments and shown as mean ± SD; each symbol represents one mouse (n = 12); Mann-Whitney test. (**I**) HA-specific IgG^+^ and IgA^+^ PCs of MHCII^+^ (MHCII sufficient), MHCII^-^ (MHCII deficient) and MHCII^B+^ (MHCII only in B cells) mice at 84 dpi. Total CD4^+^ T cells purified from uninfected C57BL/6 mice were transferred to MHCII^-^ and MHCII^B+^ mice 1 day before the IAV infection to reconstitute the CD4^+^ T cell pool. ELISPOT data are pooled from 2 independent experiments and shown as mean ± SD; each symbol represents one mouse (n = 10 for MHCII^+^, n = 9 for MHCII^-^ and MHCII^B+^); Kruskal-Wallis test with Dunn’s multiple comparison test.

We next asked whether, like RV infection, IAV-specific IgA response would require T_FH_ cells and T:B interaction. Indeed, compared to littermate controls, T_FH_-deficient Bcl6^ΔT^ mice and MHCII^ΔB^ mice both exhibited a significant reduction in the number of HA-specific IgG^+^ and IgA^+^ PCs in the NM, lung and BM at 84 dpi (Figure 7C and (S7B). These data indicate that the development of IAV-specific IgA^+^ LLPCs requires crosstalk between T_FH_ cells and Ag-presentation by B cells, as we had observed in the RV infection.

Next, we explored the role of T_FH_1 cells in the generation of IgA^+^ LLPCs. Using previously characterized PR8 NP_311-325_ NP I-A^b^ tetramers and the *Tbx21^tdT-Cre^Ifng^YFP^* double reporter mice, we detected a significant number of NP-specific tdTomato^+^CXCR3^+^CXCR5^+^PD-1^+^ T_FH_1 cells which accounted for more than 60% of NP:I-A^b+^CXCR5^+^PD-1^+^ T_FH_ cells at 14 dpi in the mediastinal LN (Figure 7D and 7E). We also observed that ∼20% of these IAV-specific T_FH_1 cells were YFP^+^ *Ifng*-expressing cells at 14 dpi (Figure 7F and 7G). These data demonstrate that T_FH_1 cells are a sizable T_FH_ cell subset generated in response to intranasal IAV infection. We next generated mixed-BM chimeras deficient in T_FH_1 cells as in Figure 5A. Similar to RV infection, the absence of T_FH_1 cells resulted in a significant reduction in HA-specific IgA^+^ LLPCs but not IgG^+^ LLPCs in the NM and lung of experimental chimeras (*Bcl6^fl/fl^Tbx21^tdT-Cre^*) relative to the control chimeras (*Bcl6^+/+^Tbx21^tdT-Cre^*) at 84 dpi (Figure 7H), although the reduction in IgA^+^ LLPCs was not as profound as what was observed for RV infection (Figure 5D). These data indicate that T_FH_1 cells are required for the optimal accumulation of IgA^+^ LLPCs in response to intranasal IAV infection.

Lastly, to assess the sufficiency of B cell Ag presentation, we took the same conditional MHCII expression approach used in the RV experiment. Unlike the RV infection scenario (Figure 3J), both HA-specific IgA^+^ as well as IgG^+^ PCs were virtually undetectable in MHCII^B+^ mice, similar to what was observed in MHCII^-^ mice (Figure 7I). These data indicate that, unlike oral RV infection, B cell Ag presentation is not sufficient for inducing IgA^+^ LLPCs in response to intranasal PR8 infection, indicating unique APC requirements in generating a virus-specific long-lived IgA response at different mucosal sites.

## Discussion

A durable and effective mucosal IgA responses is a key determinant in preventing infections against enteric viruses such as RV and poliovirus, and upper airway pathogens such as SARS-CoV-2^1–4^, however the mechanisms underlying the development of these responses remain unclear. In this study, we investigated the durability, efficacy and mechanism of generating an RV-specific IgA response. We observed that RV-specific IgA^+^ PCs in the SILP persist at least 180 days, and the IgA generated by these LLPCs is highly mutated and has the capacity to block infection in naïve mice. We uncovered a unique mechanism for the initiation of the IgA response whereby expression of MHCII on B cells is both necessary and sufficient for priming T_FH_1 cells, and through their production of IFNγ these T_FH_1 cells in turn imprint SILP-homing potential on RV-specific IgA^+^ PC that is dependent on the expression of CXCR3. In the context of the IAV response, IgA^+^ PC are partially dependent on T_FH_1 cells; however, expression of MHCII on B cells is not sufficient to support this response demonstrating tissue-specific differences between airway- vs gut-draining secondary lymphoid tissues. Taken together, our data reveal an unconventional mechanism for priming RV-specific IgA responses and implicate T_FH_1 cells in both IAV and RV IgA responses.

Despite RV being cleared by 7 dpi ((Figure S1A), a persistent GC response presumably feeds the accumulating SHM that are observed over the 180-day observation period. What source of Ag drives this GC is unknown. It is possible that fragments of VP4 and VP6 are retained on follicular dendritic cells in the PP and MLN – indeed persistent SARS-CoV-2 Spike antigen has been detected in the gut of convalescents several months post-infection concomitant with evidence of increased SHM of Spike-specific Abs^44^. In the case of RV infection, one would assume that continuous PC-flow from persistent GCs helps maintain the mucosal IgA^+^ LLPC pool, although we detected clones that were only lightly mutated, suggesting that there is a blend of early and late GC emigrants that populate the SILP. Future studies could investigate what proportion of mucosal PCs are replenished, whether the rate of replenishment varies depending on the tissue or the type of infection/vaccination, and whether this rate correlates with the quantity and quality of the LLPC pool.

In response to RV infection, Ag-presentation by B cells is necessary and sufficient for the induction of Ag-specific T_FH_1 cells, which are responsible for IgA^+^ LLPC development. How B cells induce T_FH_1 differentiation remains unclear. RV infection induces massive B cell activation independently of T cells^45^. Since activated B cells upregulate CCR6, promoting their migration to the CCL20-rich PP dome^46^, a location that is ideal for capture of intestinal Ag^47^, we hypothesize that activated B cells may prime CD4^+^ T cells in this location, as has been documented in the LN and spleen^48–50^. On the other hand, Ag-presentation by cDCs is not required for the induction of RV-specific T_FH_1 cells but is necessary for the development of an IgG response. Indeed, while deletion of T_FH_ cells prevents the formation of RV-induced IgG^+^ and IgA^+^ GC B cells, removal of T_FH_1 cells only impacts the accumulation of IgA^+^ LLPCs in the SILP, but not IgG^+^ LLPCs in the BM. This was also true for IAV-induced accumulation of IgA^+^ LLPCs in the NM and lung. Why priming by cDC vs B cells has different outcomes on T_FH_ vs T_FH_1 fate is not known. Given that CD25, ICOSL and OX40L on cDCs are critical for systemic IgG responses^51,52^, these as well as other cDC-derived molecules may skew towards T_FH_ cells that support the differentiation of IgG^+^ LLPCs.

In contrast to RV infection, Ag-presentation by B cells was not sufficient for the development of IgA^+^ LLPCs in response to IAV, indicating a requirement for other APCs in addition to B cells for the IgA response to a respiratory virus like IAV compared to an enteric virus like RV. We hypothesize that the site of priming may make a difference. For example, while PP are overlaid with M cells which take up luminal antigens and deliver them into the underlying sub-epithelial dome, M cells are very scarce in the lung at steady-state^53^. Although the nasal-associated lymphoid tissue (NALT), a potential induction site in the upper respiratory tract, has an analogous M cell-dome structure similar to the PP, NALT was recently reported to harbor an insufficient number of CD4^+^ T cells to support primary immune response to vaccination – pre-expansion of Ag-specific CD4^+^ T cells was required to visualize a response^54^. Further studies are needed to better understand the initiating mechanisms in respiratory tract-draining lymphoid tissues that ultimately support the generation of IgA^+^ LLPCs.

We found that IFNγ expression by T-bet^+^ T_FH_1 cells is required for imprinting IgA^+^ PC migration to the SILP in response to RV infection. T-bet is a master regulator for T_H_1 cells and a repressor of T_FH_ cell differentiation by diverting BCL6 away from its target genes or repressing BCL6 expression, thereby bifurcating T_H_1 vs T_FH_ cell differentiation^55–57^. However, several studies have shown that T-bet can be expressed by T_FH_ cells, although most of these studies found that its expression is transient^37,38,58,59^. Remarkably, we observed that the T-bet^+^ VP6-specific T_FH_1 cells in pooled PP and MLN for at least 8 weeks after RV infection. This is consistent with recent work from Rudensky and colleagues who observed that T-bet^+^ T_FH_1 cells are a stable population following IAV infection using a T-bet fate mapping system^38^. Only a limited number of studies have addressed the role of T_FH_1 cells in the context of immune responses: they are dispensable for IgG class switching and IgG Ab production and are crucial for the formation of tissue-resident memory B cells in the lung following IAV infection^37–39^. To our knowledge, we are the first to report a role for T_FH_1 cells in generating a long-lived virus-specific IgA response.

While B cell intrinsic IFNγR signaling was not required for the initiation of a GC response or PC induction in the PPs, it was crucial for the upregulation of CXCR3 on IgA^+^ PCs which was required for their accumulation in the SILP. This is consistent with previous observations reporting a role for IFNγR signaling in the cell-intrinsic upregulation of CXCR3 in CD4^+^ T cells and NK cells^41,42^. Where and how B cells interact with T_FH_1 cells to receive IFNγ is not known. Our scRNA-seq data revealed that RV-specific T_FH_1 cells highly express *Cd40l* and *Il21*, both of which are important for the positive selection of GC B cells into PCs^12^. Moreover, a subset of T_FH_1 cells was CD90^low^, a GC-resident phenotype^36^. Considering these observations, IgA^+^ B cells may interact with T_FH_1 cells within the GC to receive IFNγ as well as CD40L and IL-21, thereby driving both their differentiation into a PC fate as well as their expression of CXCR3. On the other hand, the RV-specific IgG response was unaffected by the absence of IFNγR on B cells. While somewhat unexpected, IFNγ from T_FH_ cells has been shown to be dispensable for IgG2c class switching in response to IAV infection^38,60^, consistent with what we observed for RV infection. It is also possible that IFNγR-deficient B cells may undergo class switching to IgG1 instead of IgG2c, thereby generating a normal overall IgG response, as recently shown in an immunization model^61^. Nevertheless, our results clearly demonstrate the critical role of T_FH_1-derived IFNγ in generating IgA^+^ LLPCs in response to RV infection.

CXCR3 is dispensable for GC development or systemic IgG production in response to immunization or viral infection^62,63^, while it is required for localization of IgA^+^ memory B cells and PCs in the lung following IAV infection^43^. Likewise, we found that B cell intrinsic CXCR3 was dispensable for the accumulation of IgG^+^ PCs in the BM but essential for the accumulation of IgA^+^ PCs in the SILP following RV infection. In response to RV, it is plausible that the expression of CXCR3 on IgA^+^ PCs works additively with CCR9 and Integrinα_4_β_7_, enabling RV-specific IgA^+^ PCs to gain a migration advantage over the large number of competitor IgA^+^ PCs which are constitutively generated in the gut. In the context of IAV infection, inflammation triggers inducible expression of CXCR3 ligands by various cell types in the respiratory tract, such as monocytes and endothelial cells^64^, which may contribute to the recruitment of IAV-specific IgA^+^ PCs to these tissues. PC survival is thought to depend on a survival niche formed by several cell types in the BM^2,13^. Given the recent study from Nutt and colleagues who observed that PCs in non-lymphoid mucosal tissues such as the gut and genital tract are extremely long-lived^65^, it is plausible that such a PC survival niche exist in the mucosae. Hence, we speculate that the CXCR3 axis guides virus-specific IgA^+^ PCs to their survival niche in these tissues.

Collectively, our data demonstrate a critical role for cDC-independent unconventional priming of T_FH_1 cells which, through an IFNγ-dependent mechanism, support the localization of CXCR3^+^IgA^+^ LLPCs in the SILP in response to RV. Because we show that IgA responses generated by this axis are durable and efficient in inducing protective immunity, our findings provide valuable insights for designing next-generation mucosal vaccines that efficiently protect against mucosal viral infections.

## Methods

### Mice

C57BL/6J, CD45.1 (B6.SJL-*Ptprc^a^Pepc^b^*/BoyJ), *Bcl6^fl/fl^* [B6.129S(FVB)-*Bcl6^tm1.1Dent^*/J, strain #023727]^25^, *Cd4^Cre^* [B6.Cg-Tg(Cd4-cre)1Cwi/BfluJ strain #022071]^66^, Prdm1-YFP [B6.Cg-Tg(Prdm1-EYFP)1Mnz/J, strain #008828]^67^, J_H_T (B6.129P2-*Igh-J^tm1Cgn^*/J, strain #002438)^68^, *H2-Ab1^fl/fl^* (B6.129X1-*H2-Ab1^b-tm1Koni^*/J, strain #013181)^69^, *Mb1^Cre^* [B6.C(Cg)-*Cd79a^tm1(cre)Reth/EhobJ^*, strain #020505]^70^, *Zbtb46^Cre^* (B6.Cg-*Zbtb46^tm3.1(cre)Mnz^*/J, strain #028538)^71^, *Cd19^Cre^* [B6.129P2(C)-*Cd19*^tm1(cre)Cgn^/J, strain #006785]^72^, *Tcrb^-/–^* (B6.129P2-*Tcrb^tm1Mom^*/J, strain #002118)^73^, *Ifng^YFP^* (B6.129S4-*Ifng^tm3.1Lky^*/J, strain #017581)^40^, *Ifng^-/–^* (B6.129S7-*Ifng^tm1Ts^*/J, strain #002287)^74^, *Ifngr1^-/–^*(B6.129S7-*Ifngr1^tm1Agt^*/J, strain #003288)^75^, *Cxcr3^-/–^*(B6.129P2-*Cxcr3^tm1Dgen^*/J, strain #005796) mice were obtained from The Jackson Laboratory. *H2-Ab1^loxP-Stop-loxP^* mice^29^ were provided by G. Wu (Washington University). *Tbx21^tdTomato-T2A-Cre^* mice^35^ were generated and provided by A. Rudensky (Memorial Sloan Kettering Cancer Center). Mice were subsequently bred in-house under specific pathogen free conditions at the University of Toronto, Division of Comparative Medicine. Mice were provided with a standard irradiated chow diet (Envigo, Teklad, 2918) and acidified water (reverse osmosis and ultraviolet sterilized) ad libitum. Female and male mice were used at 6-14 weeks of age. All animal procedures were performed ethically using age- and sex-matched littermates under animal use protocols approved by the University of Toronto Animal Care Committee.

### Bone marrow chimeras

Single cell suspensions of BM cells were prepared from tibiae and femurs as described above and 5 × 10^6^ cells were transferred intravenously into lethally irradiated (2 × 5.5 Gy, 4 hr apart) recipient mice. Recipient mice were given neomycin sulfate in their drinking water (2 g/L) for the first two weeks post-irradiation and left for an additional 6 weeks before experimentation to ensure full reconstitution of the immune system.

### Infections and treatments

For RV infection, mice were given 100 µL of 1.33% NaHCO_3_ prior to infection to neutralize the stomach acid and then inoculated with 100 µL of 10^4^ SD_50_ RV ECw (a gift from H. Greenberg, Stanford University) by oral gavage. For IAV infection, mice were anesthetized with isoflurane and inoculated intranasally with 30–40 μL of 5 × 10^4^ TCID_50_ of A/PR/8/34 H1N1 PR8 (provided by T. Watts, University of Toronto) grown in eggs. For *in vivo* CD40L blocking, mice were injected i.p. with 0.5 mg of Armenian hamster IgG (BioXCell) or anti-CD40L Ab (clone MR1, prepared in house), every other day from 0 to 14 dpi.

### Cell lines

ExpiSf9 cells were used to produce recombinant RV VP4 and VP6 and grown in ExpiSf CD medium at 27°C and 125 rpm. Expi293F cells were used to produce recombinant IgA and PR8-HA and grown in Expi293 Expression Medium (Thermo) at 37°C, 8% CO_2_ and 125 rpm. HEK293T cells were used to produce recombinant IgA and grown in D-MEM supplemented with 10% FBS and 1× Penicillin/Streptomycin at 37°C, 5% CO_2_.

### Recombinant viral proteins

Full length sequences of RV ECw VP4 and VP6 were cloned from the small intestine of C57Bl/6 mice infected with RV ECw and sequenced by Sanger sequencing as previously described^76^. The codon optimized full-length sequences of VP4 and VP6 along with a C-terminal flexible linker (GGGGSGGGGSG), an octa histidine-tag (HHHHHHHH) and an AviTag (GLNDIFEAQKIEWHE) were synthesized by GenScript into pFastBac1. Recombinant VP4 and VP6 were expressed in ExpiSf9 cells using ExpiSf Baculovirus Expression System (Thermo) as per the manufacturer’s instructions. ExpiSf9 cells were infected with recombinant baculovirus P0 or P1 stock at a multiplicity of infection (MOI) of 0.5 (VP4) or 1.0 (VP6) and harvested by centrifugation at 72-86 hr postinfection. MOI was determined by flow cytometric analysis for the expression of envelope protein gp64 on infected cells as per the manufacturer’s instructions. For purification of VP4 and VP6, recombinant baculovirus-infected ExpiSf9 cells were lysed by three times freeze/thaw cycles in NPI-0 (50 mM NaH_2_PO_4_, 500 mM NaCl, pH 8.0) supplemented with cOmplete protease inhibitor cocktail (Roche, 11836170001). The lysate was clarified by centrifugation at 20,000 ×g for 20 min, diluted with NPI-0 and incubated with TALON affinity resin (Takara) at 4°C for at least 1 hr with gentle shaking. The resin was washed with NPI-5 (NPI-0 with 5 mM imidazole) and proteins were eluted with NPI-250 (NPI-0 with 250 mM imidazole). The buffer was exchanged to PBS by dialysis and stored at -80°C until use.

For production of HA, a codon optimized sequence of HA extracellular domain from A/PR/8/34 PR8 (P03452) with a C-terminal trimeric Foldon of T4 fibritin (GYIPEAPRDGQAYVRKDGEWVLLSTFL), AviTag (GS<u>GLNDIFEAQKIEWHE</u>) and octa histidine-tag (G<u>HHHHHHHH</u>) was synthesized by GenScript into pcDNA3.4. Two cysteine residues (L20C and G373C) were introduced to stabilize HA-trimer^77^, and a Y91F substitution was introduced to prevent non-specific binding to sialic acid^78^. Recombinant HA was expressed in Expi293F cells using Expi293 Expression System (Thermo) as per the manufacturer’s instructions. For purification of HA, culture supernatants of HA-transfected Expi293F cells were collected 72-96 hr post-transfection and were clarified by centrifugation at 5,000 ×g for 20 min. The clarified supernatant was diluted with equal amount of PBS and supplemented with 1M HEPES buffer pH 7.9 to a final concentration of 20 mM and 5M NaCl to a final concentration of 300 mM and incubated with TALON affinity resin (Takara) at 4°C for at least 1 hr with gentle shaking. The resin was washed with NPI-5, and the protein was eluted with NPI-250. The buffer was exchanged to PBS using Amicon Ultra-15 centrifugal filter units.

Site-directed biotinylation was performed using the BirA enzyme (Avidity, BirA500) as per the manufacturer’s instructions. Biotinylated recombinant proteins were then mixed with fluorophore conjugated SA at a 4:1 molar ratio to generate Ag tetramer. The Ag tetramers prepared individually were then mixed and supplemented with free D-biotin (Sigma, B4639) to a final concentration of 10 µM to minimalize cross-binding of Ag tetramers. This Ag tetramer cocktail was used for cell staining followed by flow cytometry.

### ELISAs

For the detection of RV Ag shedding in feces, mouse fecal pellets were collected and homogenized using plastic rods in PBS supplemented with protease inhibitor cocktail (Millipore, 539134) (5 µL/mg stool), then the homogenates were clarified by centrifugation at 10,000 ×g for 10 min. 96-well MaxiSorp plates (Nunc, 430341) were coated with sheep anti-rotavirus Ab capture antibody (Bio-Rad, 8130-045) at 1:2000 dilution in PBS at 4°C overnight, subsequently washed and blocked with PBS with 5% skim milk at RT for 1 hr. The clarified fecal homogenates were serially diluted, added to plates and incubated at RT for 1 hr. Plates were then washed and incubated with mouse IgG2b anti-rotavirus Ab (Bio-Rad, MCA2636) at 1:200 dilution in PBS with 5% skim milk at RT for 1 hr, then washed and incubated with HRP-conjugated goat anti-mouse IgG2b Ab (SouthernBiotech, 1090-05) at 1:2000 dilution in PBS with 5% skim milk at RT for 1 hr, washed then developed with TMB Solution (Thermo, 00-4201-56) followed by addition of 1M H_2_SO_4_ to stop the reaction. Absorbance at 450 nm was then measured.

For the detection of Ab binding to VP4 and VP6, 96-well MaxiSorp plates (Nunc, 430341) were coated with 10 µg/mL of recombinant VP4 or VP6 at 4°C overnight. Plates was washed and blocked with PBS with 3% BSA at RT for 1 hr. Serially diluted samples were loaded onto plates and incubated at RT for 1 hr. Plates were washed and incubated with HRP-conjugated anti-mouse IgA Ab (SouthernBiotech, 1040-05) at 1:2000 dilution in PBS with 0.3% BSA. Plates were washed and developed with TMB Solution (Thermo, 00-4201-56), followed by addition of 1M H_2_SO_4_, then absorbance at 450 nm was measured.

For the measurement of IgA concentration, 96-well MaxiSorp plates (Nunc, 430341) were coated with goat anti-mouse IgA (SouthernBiotech, 1040-01) at 1:1000 dilution in PBS at 4°C overnight. Plates were washed, blocked with PBS with 3% BSA at RT for 1 hr. Serially diluted samples were loaded onto plates and incubated at RT for 1 hr. Plates were then washed and incubated with HRP-conjugated anti-mouse IgA Ab (SouthernBiotech, 1040-05) at 1:2000 dilution in PBS with 0.3% BSA, washed and developed with TMB Solution (Thermo, 00-4201-56), followed by addition of 1M H_2_SO_4_. Absorbance at 450 nm was measured. PBS supplemented with 0.05% Tween20 (PBS-T) was used for all washing steps in ELISA.

### Preparation of single cell suspensions

The PP, MLN and mediastinal LN were collected and processed with the frosted ends of microscope glass slides. BM cells were collected by centrifugation of punctured tibiae and femurs at up to 10,000 ×g for 10 sec and then treated with ACK buffer (155 mM NH_4_Cl, 10 mM KHCO_3_, 0.1 mM EDTA, pH 7.3) to lyse red blood cells (RBCs). Blood was collected from the saphenous vein of mice using Microvette capillary tubes (Sarstedt, 16.443.100). RBCs were then lysed with ACK buffer.

For the isolation of SILP, PPs and the luminal contents were removed from the SI. The SI was cut longitudinally, then cut into 1-cm pieces and washed with PBS. The tissues were incubated at 37°C for 40 min in HBSS supplemented with 3% FBS, 10 mM HEPES pH 7.5 and 5 mM EDTA with continuous agitation. The tissues were washed three times with PBS and incubated at 37°C for 30 min in RPMI 1640 supplemented with 3% FBS, 10 mM HEPES pH 7.5, 250 µg/mL collagenase IV (Sigma, C5138) and 50 µg/mL DNase I (Roche, 11088866001). The remaining tissues were further digested by another 30-min incubation in a fresh digestion medium. After enzymatic digestion, the homogenates were filtered through a 70-µm cell strainer and then lymphocytes were enriched by centrifugation at 800 ×g for 10 min in 40% Percoll at RT.

To collect the nasal mucosa (NM), mice were euthanized and perfused transcardially with 10 mL of ice-cold PBS. The lower jaw and tongue, eyes, skin and connective tissues were removed from the head. The upper plate, to which the nasal associated lymphoid tissue (NALT) is attached, was removed with fine scissors and forceps as previously described^54^, to avoid contamination of NALT. The entire skull and nose were cut into two along the midline using a fine razor blade and then the brain was removed. Three regions of the nasal tissues – the respiratory mucosa, the olfactory mucosa and the lateral nasal gland – were collected, pooled and were referred to collectively as the NM. The NM was then chopped into small pieces and digested at 37°C for 30 min in RPMI 1640 supplemented with 3% FBS, 10 mM HEPES pH 7.5, 1 mg/mL collagenase D (Roche, 11088866001) and 100 µg/mL DNase I (Roche, 11088866001). The homogenates were filtered through a 70-µm cell strainer and remaining tissues were dissociated using a 3-mL syringe plunger.

All lobes of the lung were collected after perfusion with 10 mL of ice-cold PBS and were chopped into small pieces then digested at 37°C for 45 min in RPMI 1640 supplemented with 3% FBS, 10 mM HEPES pH 7.5, 1 mg/mL collagenase D (Roche, 11088866001) and 100 µg/mL DNase I (Roche, 11088866001). The homogenates were filtered through a 70-µm cell strainer and remaining tissues were dissociated using a 3-mL syringe plunger. Cells were then treated with ACK buffer to lyse RBCs.

All cell suspensions were washed using HBSS with 3% FBS and filtered through a nylon mesh to remove debris prior to downstream analysis.

### Flow cytometry and cell sorting

Single-cell suspensions were incubated with 2 µg/mL of anti-CD16/32 Ab (2.4G2, BioXCell, BE0307) for at least 10 min at 4°C to block Fc receptors and stained for cell surface markers and viability dye on ice for 25 min in FACS buffer (PBS with 0.5 % FBS and 2 mM EDTA). Cells were washed twice with FACS buffer. For intracellular staining, cells stained for cell surface markers and viability dye were washed with PBS and fixed with the Foxp3/Transcription factor staining kit (Thermo, 00-5523-00) as per the manufacturer’s instructions. Cells were washed with 1× permeabilization buffer and stained for transcription factors on ice overnight or for immunoglobulin isotypes at RT for 30 min. Cells were washed twice with 1× permeabilization buffer and once with FACS buffer. The staining antibodies for flow cytometry were purchased from Thermo, Biolegend, BD Biosciences and SouthernBiotech. The following reagents were used for staining: B220 (RA3-6B2), BCL6 (7D1), CD11b (M1/70), CD11c (N418), CD138 (281-2), CD3 (17A2), CD38 (90), CD4 (RM4-5), CD40L (MR1), CD44 (IM7), CD45 (30-F11), CD45.1 (A20), CD45.2 (104), CD69 (H1.2F), CD90.2 (30-H12), CD98 (RL388), CXCR3 (CXCR3-173), CXCR5 (2G8), F4/80 (BM8), GL7 (GL7), IgA (goat polyclonal), IgD (11-26c.2a), IgG1 (RMG1-1), IgG2b (RMG2b-1), IgG2c (RMG2a-62), IgG3 (goat polyclonal), IgM (RMM-1), IRF4 (3E4), Ki-67 (SolA15), Ly6A (D7), MHCII (M5/114.15.2), PD-1 (J43), PSGL-1 (2PH1), T-bet (4B10), TACI (ebio8F10-3), TIGIT (1G9) and Streptavidin. For IgG staining, a cocktail of anti-IgG1, anti-IgG2b, anti-IgG2c and anti-IgG3 was used. For Ag (B cell) tetramer staining, up to 3 × 10^7^ cells were prepared in V-bottom 96-well plate and stained with 50 µL of Ag tetramer cocktail as described above containing 180 ng of VP4 (2 color, total 360 ng) and/or 90 ng of VP6 (2 color, total 180 ng) on ice for 1 hr. Finally, samples were filtered through a 40-µm strainer and acquired on BD LSR X20, FACSymphony A3 or A5 analyzer or sorted on FACSAria III using FACSDiva software. The absolute number of cells were calculated using ContBright Plus Absolute Counting Beads (Thermo) as per the manufacturer’s instructions. Data were analyzed using FlowJo v10 software. For (Figure S4L, FCS files from FACSDiva software were imported into R v4.3.2 using the Bioconductor packages CytoML v2.14.0 and flowWorkspace v4.14.3 and t-SNE projection was computed using the Bioconductor package CATALYST v1.26.1. Histogram and t-SNE plots were generated using the CRAN packages ggplot2 v3.5.1, scico v1.5.0 and cowplot v1.1.3.

### ELISPOTs

MultiScreen plates (Milipore, MSHAS4510 and MSIPS4W10) were coated with 10 µg/mL of recombinant VP4, VP6 or PR8-HA at 4°C overnight. Plates were washed three times with PBS and blocked with complete media (RPMI 1640 containing 10% FBS, 10 mM HEPES pH 7.5, 1 mM sodium pyruvate, 1× MEM non-essential amino acid, 1× GlutaMAX, 1× Penicillin/Streptomycin and 55 µM 2-mercaptoethanol) at RT for 1 hr. The plate was loaded with serially diluted single cell suspensions in complete media and incubated at 37°C, 5% CO_2_ for 16-24 hr. The plate was then washed with PBS-T and incubated with HRP-conjugated anti-mouse IgA Ab (SouthernBiotech, 1040-05) or HRP-conjugated anti-mouse IgG Ab (SouthernBiotech, 1030-05) at 1:1000 dilution in PBS with 0.3% BSA at RT for 1 hr. Then, plates were washed and developed with AEC substrate (Vector Laboratories, SK-4200).

### Antibody sequencing, analysis and cloning

V(D)J regions of *Igh*, *Igk* and *Igl* genes of single IgA^+^ PCs were amplified by nested-PCR using V gene-specific forward primers and C region-specific reverse primers as previously described^79^. PCR products were processed using ExoSAP-IT PCR product cleanup reagent (Thermo, 78201) and sequenced by Sanger sequencing. Sequence analysis was performed using IMGT/HighV-QUEST and in-house pipeline to determine the V(D)J rearrangement and the number of SHM compared to putative germline sequences. Sequences that shared V_H_/J_H_ genes, had the same CDR3 lengths, and contained at least 75% CDR3 nucleotide identity were grouped and classified into clonal lineages. Only cells productively rearranged V(D)J were used for V_H_ + V_L_ or V_H_ mutation analyses. GCtree^80^ was used to infer clonal lineage trees and the unmutated VDJ sequences were used for outgroup rooting.

For construction of mouse IgA expression vectors, a mouse IgA sequence with a 5’ AfeI restriction site, which was introduced by silent mutations, was synthesized by GenScript and inserted into SalI-HindIII site of human IgG1 expression vector (a gift from M. Nussenzweig, the Rockefeller University)^79^ to replace the human IgG1 sequence. V(D)J regions of *Igh* and *Igk* genes were amplified from the first PCR product of the nested-PCR as previously described^79^. V_H_-, V_K_- and J_K_-specific primers^79^ and J_H_-specific primers containing an AfeI site as listed below were used for this cloning-PCR.

T4mRA_I: ATGGTGGGATTTCTAGCGCTCTCTGAGGAGACGGTGACCGTGG
T4mRA_II: ATGGTGGGATTTCTAGCGCTCTCTGAGGAGACTGTGAGAGTGG
T4mRA_III: ATGGTGGGATTTCTAGCGCTCTCTGAGGAGACAGTGACCAGAG
T4mRA_IV: ATGGTGGGATTTCTAGCGCTCTCTGAGGAGACGGTGACTGAGG

The AgeI-AfeI linearized mouse IgA expression vector and unpurified cloning-PCR product were used as templates for NEBuilder HiFi DNA assembly (New England Biolabs), and assembled DNA was transformed in DH5α competent bacteria. All plasmid sequences were verified by Sanger sequencing.

### Recombinant IgA production

To produce dimeric recombinant IgA, codon optimized mouse IgJ sequence was synthesized by Twist Bioscience into the NotI-NheI site of the pTwist CMV BetaGlobin vector. This IgJ sequence was also cloned into the BamHI-NotI site of the pMXs-IRES-EGFP retroviral vector. HEK293T cells were retrovirally transduced with pMXs-IgJ-IRES-EGFP and the stable transformants were sorted based on the EGFP expression by flow cytometry. For *in vitro* validation of rIgA, paired mouse IgA and IgK expression vectors were co-transfected in HEK293T-IgJ cells using PEI Max (Mw 40,000) (Polysciences, 24765). Culture supernatants were collected 72 hr after transfection and used for ELISA as described above. For *in vivo* administration, rIgAs were expressed by co-transfection of paired mouse IgA and IgK expression vectors and pTwist CMV BetaGlobin-IgJ in Expi293F cells at a 1:1:1 ratio using Expi293 Expression System (Thermo) as per the manufacturer’s instructions. For purification of rIgA, culture supernatants of rIgA-transfected Expi293F cells were collected 96-120 hr post-transfection and clarified by centrifugation at 5,000 ×g for 20 min. The clarified supernatants were diluted with equal amount of PBS and incubated with Capto L resin (cytiva) at 4°C for at least 1 hr with gentle shaking. For rIgAs unable to bind to protein L, VP4- or VP6-immobilized resins were made by conjugating recombinant VP4 or VP6 with NHS-activated Sepharose 4 FF (Cytiva) as per the manufacturer’s instructions and used for affinity purification. After the incubation with culture supernatant, the resin was washed with PBS, and rIgAs were eluted with 0.1 M Glycine-HCl pH 2.9, then 1M Tris-HCl pH8.0 was added to neutralize pH. The buffer was exchanged to PBS using Amicon Ultra-15 centrifugal filter units.

### Peptide library and AIM assay

Overlapping peptide library was synthesized by GenScript as 15-mers spanning the entire sequences of VP4 or VP6, overlapping by 11-mer. Total 191 peptides for VP4 and 97 peptides for VP6 were synthesized. Individual peptides were resuspended in DMSO, pooled and stored at – 80°C until use. For the AIM assay, single-cell suspensions were prepared from pooled PPs and MLNs of B6 mice at 7 dpi with RV and were seeded at 2 × 10^6^ cells per well of flat-bottom 96-well plates. Cells were stimulated with 1 µg/mL of peptide pools (concentration of each peptide), 5 µg/mL of single peptides or 5 µg/mL of Concanavalin A (ConA) in complete media in the presence of 2 µg/mL of PE-conjugated anti-CD40L for 18 hr at 37°C, 5% CO2. After incubation, cells were stained for cell surface markers and viability dye at RT for 20 min in FACS buffer and analyzed by flow cytometry as described above.

### MHCII tetramer production

VP4 and VP6 peptide:I-A^b^ tetramers were generated by an in-house peptide exchange method. The nucleotide sequence encoding the murine CLIP_88–99_ followed by a linker with an internal thrombin cleavage site (SQ<u>MRMATPLLM</u>RGGGGS<u>LVPRGS</u>GGGG - nonamer binding core of the CLIP peptide and thrombin cleavage sites are underlined) were cloned into an expression plasmid^32^ upstream of the nucleotide sequence encoding the I-A^b^ β extracellular domain linked to a basic leucine zipper and hexa histidine tag. This plasmid was co-transfected into S2 insect cells with a plasmid encoding the I-A^b^ α chain extracellular domain linked to an acidic leucine zipper and a BirA tag, a plasmid encoding the BirA ligase (p18-BirA) and a plasmid encoding the blasticidin resistance gene (pCo.Blast). Biotinylated CLIP(T):I-A^b^ β chain/I-A^b^ α chain monomers (CLIP(T):I-A^b^) were produced and purified from culture supernatants as previously described^81^. Monomers were tetramerized with PE-conjugated SA. The CLIP(T):I-A^b^/SA-PE tetramer was mixed with candidate VP4 or VP6 peptides individually in a 0.2 ml PCR tube such that the final concentration of tetramer and peptide was 1 µM and 100 µM, respectively. 0.1 U of thrombin (Millipore, restriction grade) were added and the mixture was incubated at 37°C for 6 hours. Five volume of 1 µM recombinant mouse H2-DM (produced in-house) in sodium citrate buffer (pH 5.4) was added to one volume of the mixture of CLIP(T)-I-A^b^ and peptide, and incubated at 37°C overnight. The mixture was then stored at 4°C until use. The sequences of candidate VP4 and VP6 peptides are listed below. VP4-p51: HSDFYIIPWAQQSLC, VP4-p52: YIIPWAQQSLCTRYI, VP4-p110: SITRTRVSGLYGLPA, VP4-p120: DDYQTPIMNSVTVRQ, VP6-p58: AERFSFPRVINSADG, VP6-p61: ADGATTWYFNPVILR, VP6-p62: TTWYFNPVILRPNNV. For the covalently linked APC-labeled VP6:I-A^b^ tetramers, the selected VP6 peptide was adapted into the affinity enhanced 4E tetramer platform as described previously^81^.

### MHCII tetramer staining

APC-labeled NP_311-325_:I-A^b^ tetramer was obtained from the National Institutes of Health (NIH) tetramer core facility. For I-A^b^ tetramer staining, PPs were incubated at 37°C for 30 min in HBSS supplemented with 3% FBS, 10 mM HEPES pH 7.5, 5 mM EDTA and 1 mM dithiothreitol with continuous agitation to remove epithelial cells and mucus. After the incubation, PPs and MLN from the same mouse were pooled and dissociated using a 3-mL syringe plunger through a 70-µm cell strainer and washed twice with FACS buffer. Mediastinal LN of PR8 infected mice were dissociated with the frosted ends of microscope glass slides. Cells were resuspended in FACS buffer containing 50 nM dasatinib (Sigma) and 2 µg/mL of anti-CD16/32 Ab (2.4G2, BioXCell, BE0307) and incubated at 37°C for 15 min. APC-labeled peptide:I-A^b^ tetramers were added to the cell suspension at 10 nM final concentration and incubated at RT for 1 hr. After washing twice with FACS buffer, cells bound to peptide:I-A^b^ tetramers were magnetically enriched using EasySep APC positive selection kit II (STEMCELL Technologies) as previously described^32^. The bead-bound cells were stained for cell surface markers, viability dye and/or intracellular transcription factors and analyzed by flow cytometry.

### CD4^+^ T cell isolation and transfer

CD4^+^ T cells were purified from spleens of naive mice by magnetic negative sorting. Single-cell suspensions of splenocytes were incubated with 2 µg/mL of anti-CD16/32 Ab (2.4G2, BioXCell, BE0307) for 15 min on ice and incubated with a cocktail of biotinylated Abs for Ter119, B220, CD138, MHCII, IgD, IgM, CD49b, CD8, CD11b, NK1.1, Ly6G, F4/80 and γδTCR on ice for 20 min. Cells were washed twice with FACS buffer and incubated with Streptavidin Particles Plus DM (BD Biosciences) (80 µL per spleen) on ice for 20 min. The unbound fraction to the beads were negatively purified using EasySep Magnet (STEMCELL Technologies), then the unbound fraction was further purified using MidiMACS Separator (Miltenyi Biotec), yielding CD3^+^CD4^+^ T cells of > 95% purity. Purified CD4^+^ T cells were washed three times with PBS and were intravenously transferred to recipient mice (2 × 10^7^ cells/mouse), 24-36 hours before infection.

### scRNA-Seq library preparation and analysis

The PPs and MLNs from RV infected mice were dissociated using a 3-mL syringe plunger through a 70-µm cell strainer and washed twice with FACS buffer. Cells from each organ were split into 2 samples and were resuspended in FACS buffer containing 2 µg/mL of anti-CD16/32 Ab (2.4G2, BioXCell, BE0307) and 20 nM of PE-labeled either VP4:I-A^b^ or VP6:I-A^b^ tetramers and incubated at RT for 1 hr. After washing twice with FACS buffer, tetramer^+^ cells were magnetically enriched using EasySep PE positive selection kit II (STEMCELL Technologies) as previously described^32,81^. The unbound fractions were also collected and pooled into one tube. The bead-bound and unbound cells were stained for cell surface markers for flow cytometric sorting as well as hashtag oligonucleotides (HTOs) to pool different samples into the same sequencing run. Each sample was stained with one unique HTO (Biolegend, TotalSeq C anti-mouse Hashtag Abs): HTO1 (M1/42; 30-F11, 155861), HTO2 (M1/42; 30-F11, 155863), HTO3 (M1/42; 30-F11, 155865), HTO4 (M1/42; 30-F11, 155867) and HTO5 (M1/42; 30-F11, 155869). Then, by flow cytometric sorting, Dump (B220, CD11b, CD11c, F4/80 and viability dye)-negative CD3^+^CD4^+^CD8^-^p:I-A^b+^ cells from the bead-bound fractions (referred as VP4:I-A^b+^ and VP6:I-A^b+^) and an equal number of Dump-negative CD3^+^CD4^+^CD8^-^p:I-A^b–^ cells from the bead-unbound fraction (referred as Total CD4^+^) were sorted into the same collection tube. The sorted cells were counted for viability and immediately subjected to library preparation using 10X Chromium Next GEM Single Cell 5’ Reagent Kits V2 and Chromium Single Cell Mouse TCR Amplification Kit according to the manufacturer’s instructions, and the scRNA-seq and TCR-seq libraries were sequenced on the Illumina NovaSeq 6000 platform at the Princess Margaret Genomics Centre in Toronto, Ontario, Canada. Two independent experiments sequencing VP4/VP6^+^ CD4^+^T cells were performed processed for each sample except for Total CD4^+^ (n = 1). Each biological replicates represents a pool of PPs and MLNs cells from > 40 mice. Raw FASTQ files from the 10X libraries were processed with CellRanger v7.1.0 using the mm10 reference genome for sample demultiplexing, barcode processing and single-cell counting. Read matrices were analyzed using the Seurat package v5.0.1 in R v4.3.2. Specifically, CellRanger output was imported using the Read10X function. Quality control was performed by excluding cells with fewer than 300 genes detected, more than 6,000 genes detected or more than 4% mitochondrial reads. Cells expressing either zero or more than one hashtag were removed from analysis. Read counts were then normalized using the NormalizeData function (default parameters). The ScaleData function was used for data scaling, regressing out the difference between the G2M and S phase scores to maintain signals separating non-cycling cells and cycling cells but to remove differences in cell cycle phase among proliferating cells out of the data. Individual samples were then integrated using the IntegrateLayers function and CCAIntegration method with default parameters. Cells were clustered using the FindNeighbours with the 30 (Figure 4B) or 10 (Figure 4D) largest principal components and the FindClusters functions with the resolution parameter equal to 0.8 (Figure 4B) or 0.5 (Figure 4D). Dimension reduction and visualization were performed with uniform manifold approximation and projection (UMAP). Differentially expressed genes were identified in Seurat with the FindMarkers function by running a Wilcoxon rank-sum test for clusters 5 and 6, and visualized using EnhancedVolcano v1.20.0. Gene set enrichment analysis (GSEA) was conducted using the package clusterProfiler v4.10.0. All visualizations of scRNA-seq data were generated using the Seurat package as well as ggplot2 v3.4.4.

### Quantification and statistical analysis

Statistical information, including n (number of mice or cells per group), number of replicates, mean and statistical significance values, are indicated in the figure legends. All statistical tests, including normality tests, were performed using GraphPad Prism 8 software. Only p values less than 0.05 were considered significant. Significance was indicated as follows: ^∗^p < 0.05, ^∗∗^p < 0.01, ^∗∗∗^p < 0.001, ^∗∗∗∗^p < 0.0001, NS: not significant. No specific method was used to determine whether the data met assumptions of the statistical approach.

## Acknowledgments

We thank A. Rudensky (Memorial Sloan Kettering Cancer Center) for *Tbx21^tdTomato-T2A-Cre^* mice, H. Greenberg (Stanford University) for RV ECw strain, M. Nussenzweig (the Rockefeller University) for Ab expression vector, F. Saori (UHN) for critical suggestion and discussion, M. Jenkins (University of Minnesota) for helpful discussion, A. Nguyen and M. Zuo (University of Toronto) for technical advice and support, C. Li (CUHK) for technical advice and assistance for RV infection, L. Ward (University of Toronto) for preparing anti-CD40L Ab, Birinder Ghumman (University of Toronto) for preparing and tittering IAV, K. Yeung (University of Toronto) for technical advice on IAV infection, the veterinary staff at DCM and the staff at the Faculty of Medicine Flow Cytometry core facility (University of Toronto) for support, and all members of the Gommerman Laboratory for help. We are grateful for funding support from the Canadian Institutes for Health Research / Instituts de recherche en santé du Canada (CIHR/IRSC) to J.L.G (FDN-159922), T.H.W (PJT-178020) and the Uehara Memorial Foundation to K.H.

## Figures

**Figure S1.**
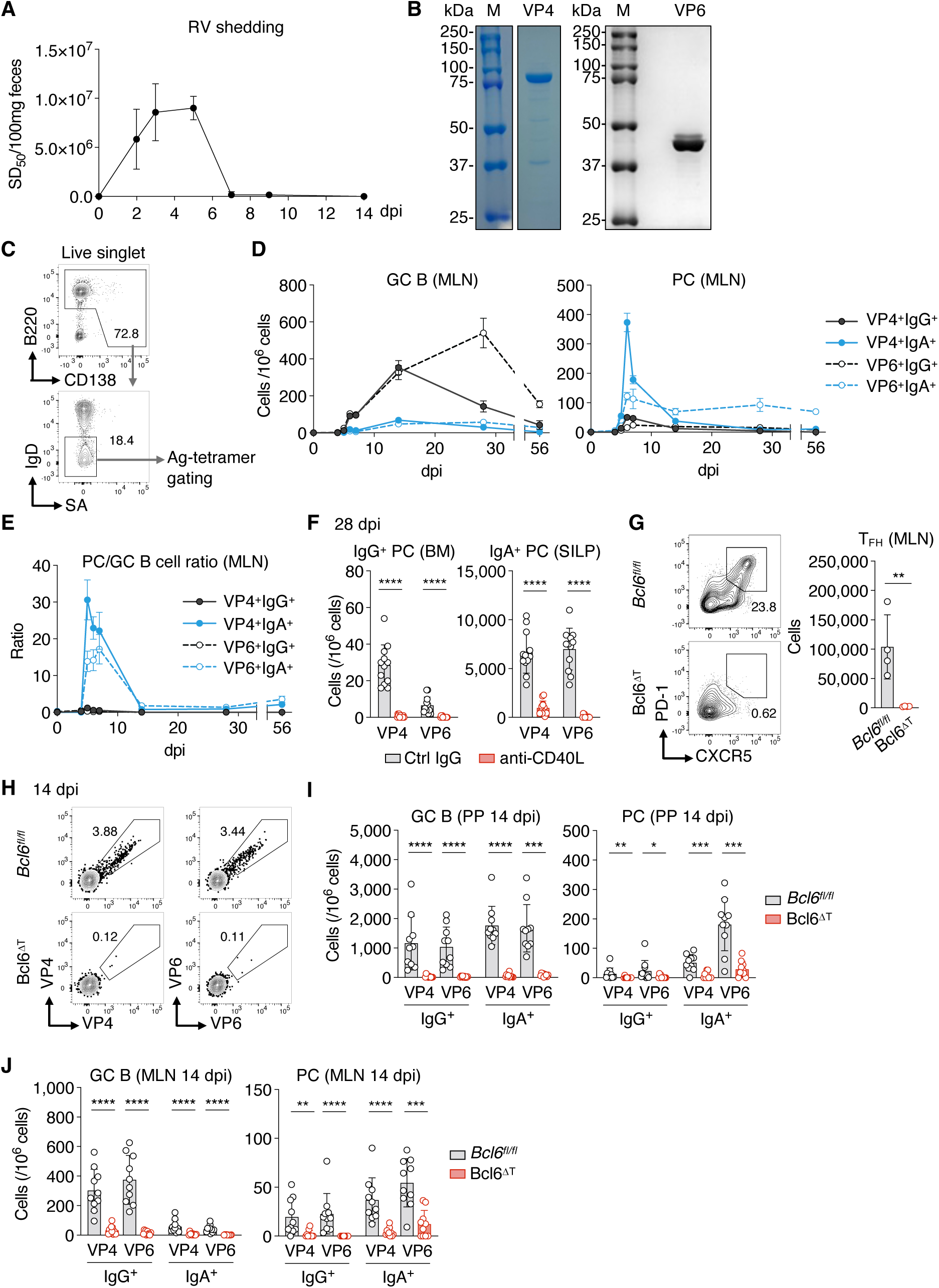
Analysis of Ag-specific GC B cells and PCs following RV infection. (**A**) RV shedding in C57BL/6 mice infected with RV measured by ELISA. Data are mean ± SD (n = 4). (**B**) SDS-PAGE of purified recombinant VP4 and VP6. SDS-PAGE gels under reducing condition were stained by Coomassie brilliant blue. Both strips (left) were cropped from the same gel. M: molecular weight marker. (**C**) Representative flow cytometry profiles showing the strategy for subsequent Ag-tetramer gating in Figure 1A. (**D and E**) Flow cytometric analysis of GC B cells and PCs in MLN after RV infection. Graph showing the number of GC B cells and PCs (D) and the ratio of PCs to GC B cells (E). Data are pooled from at least 2 independent experiments and shown as mean ± SEM (n = 4-21). Gating strategy is shown in Figure 1A. (**F**) ELISPOT assay analyzing the number of VP4- and VP6-specific IgG^+^ and IgA^+^ PCs at 28 dpi with RV in mice treated with either control Armenian hamster (Ctrl) IgG or anti-CD40L Ab. 0.5 mg of Ab was administered i.p. every other day from 0 to 14 dpi. Data are pooled from 2 independent experiments and shown as mean ± SD; each symbol represents one mouse (n = 12); Mann-Whitney test. (**G**) Flow cytometric analysis of T_FH_ cells in MLN of *Bcl6^fl/fl^* and Bcl6^ΔT^ mice at 7 dpi with RV. Representative plots showing the percentage of CXCR5^+^PD-1^+^ T_FH_ cells among CD3^+^CD4^+^CD44^hi^ cells and the graph showing the absolute number of T_FH_ cells. Data are shown as mean ± SD; each symbol represents one mouse (n = 4); Student’s t-test. (**H-J**) Flow cytometric analysis of *Bcl6^fl/fl^* and Bcl6^ΔT^ mice at 14 dpi. Representative plots showing the percentage of VP4^+^ and VP6^+^ cells among B cells and PCs in the PPs (H), graph showing the number of GC B cells and PCs in the PPs (I) and graph showing the number of GC B cells and PCs in the MLN (J). Data are pooled from 2 independent experiments and shown as mean ± SD; each symbol represents one mouse (n = 10); Mann-Whitney test. Gating strategy is shown in Figure 1A.

**Figure S2.**
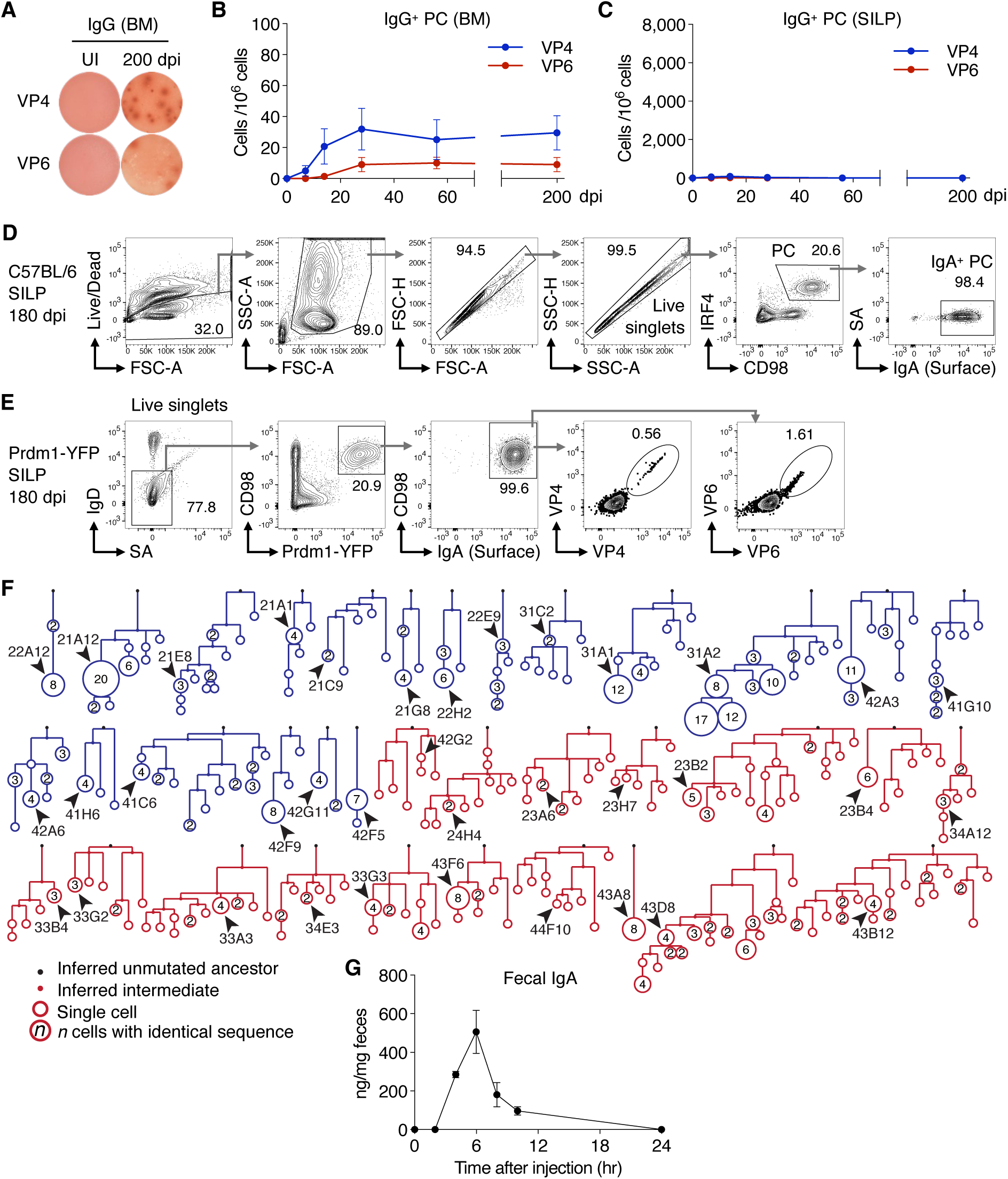
Analysis of Ag-specific PCs following RV infection. (**A-C**) ELISPOT assay detecting VP4- and VP6-specific IgG^+^ PCs after RV infection. Representative ELISPOT image (A). The number of IgG^+^ PCs in the BM (B) and the SILP (C). Data are pooled from at least 2 independent experiments and shown as mean ± SD (n = 8). (**D**) Representative flow cytometric profiles showing the gating strategy of IgA^+^ PCs in the SILP of C57BL/6 WT mice at 180 dpi. Streptavidin (SA) was used to gate out cells that non-specifically bind to SA; cell surface IgA was stained. (**E**) Representative flow cytometric profiles showing the gating strategy of IgA^+^ PCs in the SILP of *Prdm1*-YFP mice at 180 dpi with RV. (**F**) Phylogeny inferences based on *Ighv* sequences of 19 clones of anti-VP4 (blue) and 17 clones of anti-VP6 (red). Each arrowhead indicates selected cell for rIgA production used for ELISA in Figure 2I. (**G**) Time course of rIgA secretion in feces of J_H_T mice after i.v. injection of 0.5 mg rIgA. Data are mean ± SD (n = 4).

**Figure S3.**
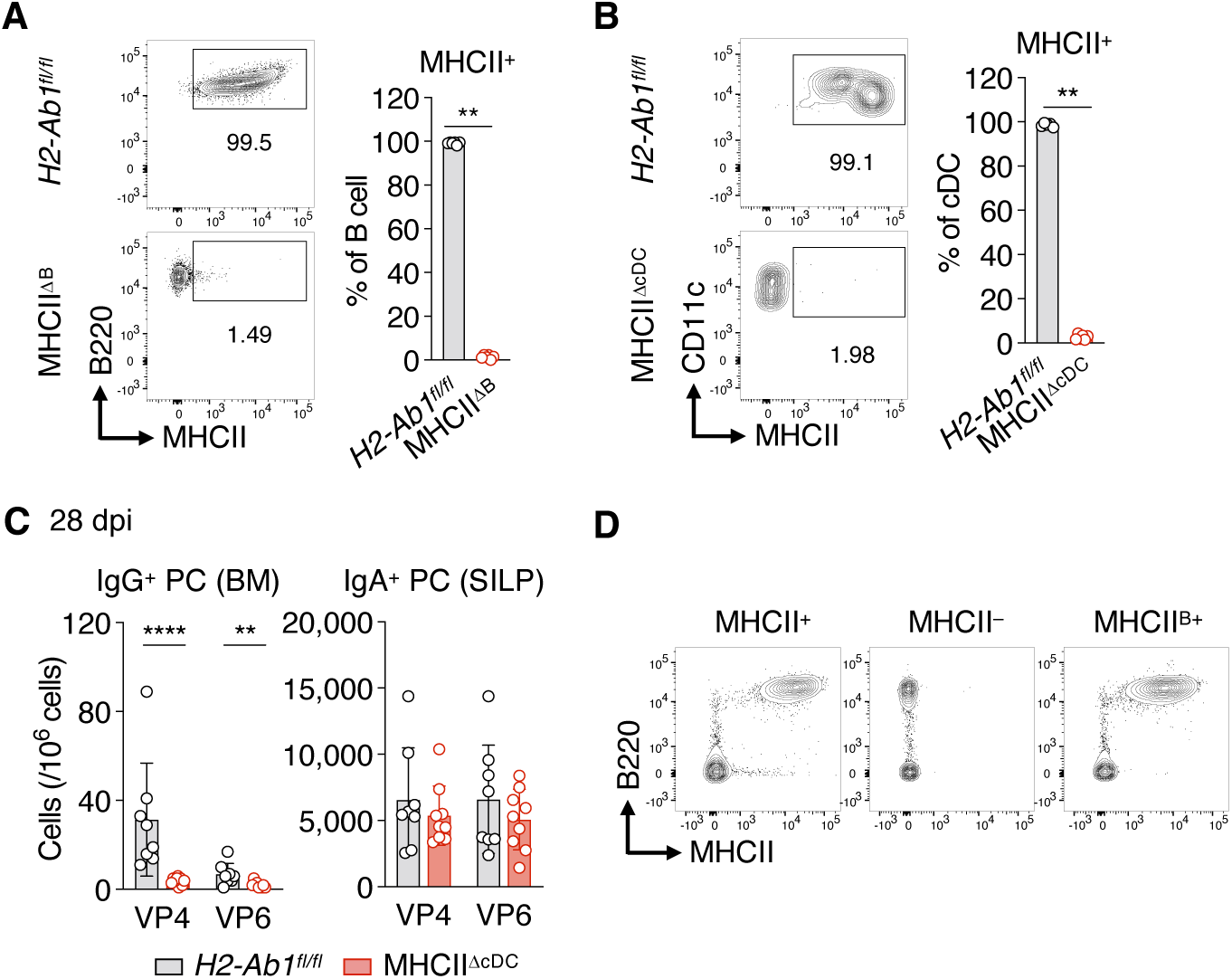
Analysis of MHCII^ΔB^, MHCII^ΔcDC^ and MHCII^B+^ mice. (**A**) Flow cytometry analyzing the percentage of MHCII^+^ cells among B cells (CD3^-^Ly6C^-^Ly6G^-^ CD11c^-^B220^+^) of *H2-Ab1^fl/fl^* and MHCII^ΔB^ mice. Data are shown as mean ± SD; each symbol represents one mouse (n = 6); Mann-Whitney test. (**B**) Flow cytometry analyzing the percentage of MHCII^+^ cells among cDCs (CD3^-^Ly6C^-^Ly6G^-^ CD11c^+^B220^-^) of *H2-Ab1^fl/fl^* and MHCII^ΔcDC^ mice. Data are shown as mean ± SD; each symbol represents one mouse (n = 5 for *H2-Ab1^fl/fl^*, n = 6 for MHCII^ΔcDC^); Mann-Whitney test. (**C**) The number of VP4- and VP6-specific IgG^+^ and IgA^+^ PCs of *H2-Ab1^fl/fl^* and MHCII^ΔcDC^ mice at 28 dpi. ELISPOT data are pooled from 2 independent experiments and shown as mean ± SD; each symbol represents one mouse (n = 8 for *H2-Ab1^fl/fl^*, n = 9 for MHCII^ΔcDC^); Mann-Whitney test. (**D**) Representative flow cytometry plots showing MHCII and B220 profiles in peripheral blood lymphocytes of MHCII^+^, MHCII^-^ and MHCII^B+^ mice.

**Figure S4.**
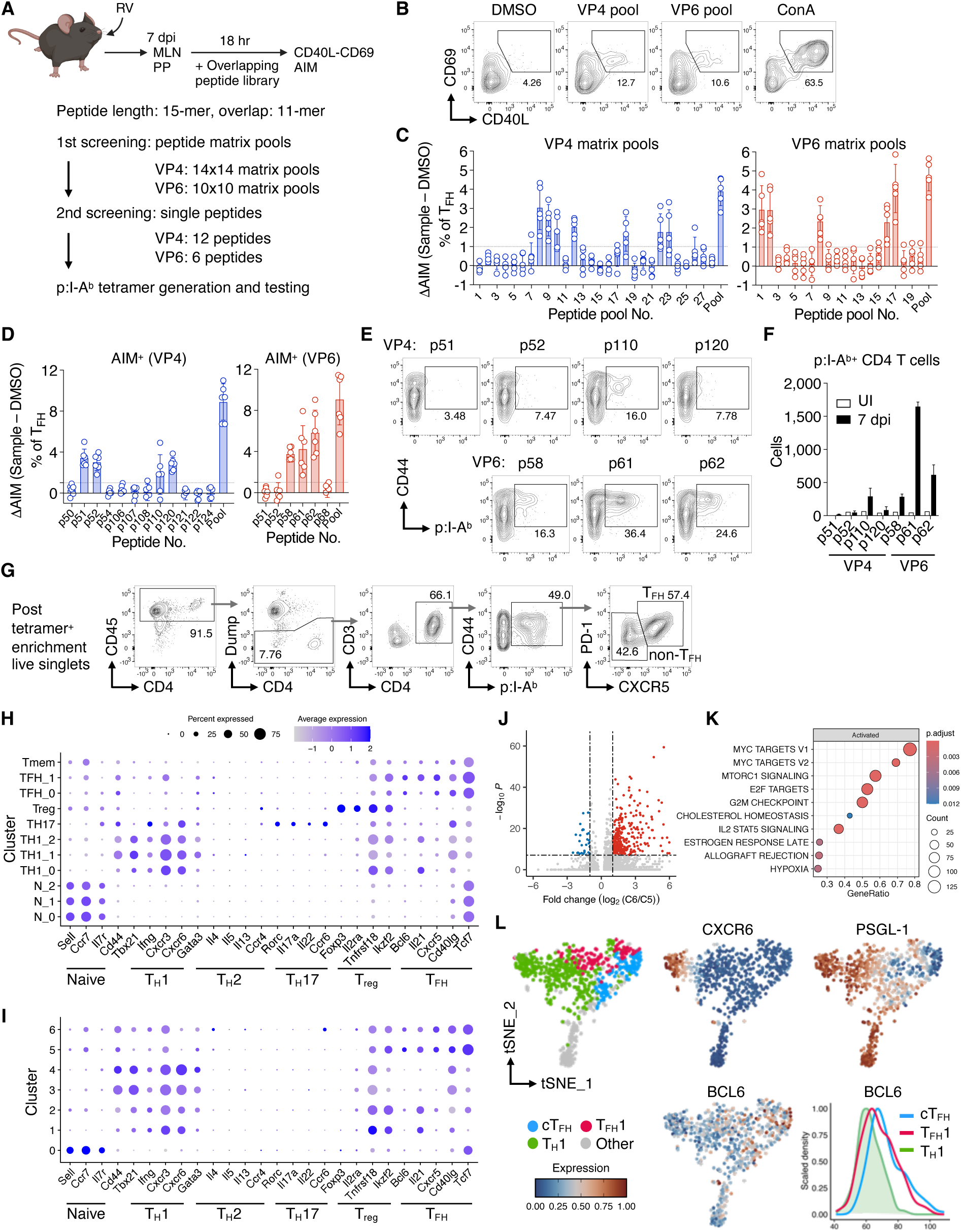
Identification and characterization of RV-specific CD4^+^ T cells. (**A-D**) T_FH_ peptide epitope mapping of VP4 and VP6 proteins. Schematic representation of the strategy (A). Representative flow cytometry plots showing the percentage of CD40L^+^CD69^+^ (activation induced marker positive, AIM^+^) cells among CD45^+^CD3^+^CD4^+^CXCR5^+^PD-1^+^ T_FH_ cells (B). Graphs showing the normalized AIM positivity (ΔAIM: %AIM^+^ in sample minus that in DMSO control) using matrix pools (C) and single peptides (D). ConA, Concanavalin A. Data are pooled from 5 (C) or 6 (D) independent experiments; each symbol represents a result of one experiment. (**E and F**) Flow cytometric analysis of peptide:I-A^b^ (p:I-A^b^) staining of PP and MLN cells from C57BL/6 mice at 7 dpi. Representative plots showing the frequencies of p:I-A^b+^ cells among CD45^+^B220^-^CD11b^-^CD11c^-^F4/80^-^CD3^+^CD4^+^ T cells (E) and graph showing the absolute number of pI-A^b+^ CD4^+^ T cells (F). pI-A^b+^ cells were enriched from pooled PP and MLN cells by magnetic sorting and used for analysis. Data are pooled from 2 independent experiments and shown as mean ± SD. UI, uninfected. (**G**) Representative flow cytometry profiles showing the gating strategy of RV-specific CD4^+^ T cells. pI-A^b+^ cells were enriched from pooled PP and MLN cells by magnetic sorting and used for flow cytometry. Dump: B220, CD11b, CD11c and F4/80. (**H**) Dot plot of scRNA-seq data of total, VP4:I-A^b+^ and VP6:I-A^b+^ CD4^+^ T cells showing expression of CD4^+^ T cell lineage-associated genes. (**I-K**) scRNA-seq analysis of VP4:I-A^b+^ and VP6:I-A^b+^ CD4^+^ T cells only. Dot plot of CD4^+^ T cell lineage-associated gene expression (I). Volcano plot of differentially expressed genes between C5 and C6 (J). Gene set enrichment analysis of differentially expressed genes between C5 and C6 (K). Cells are pooled from 2 independent experiments. (**L**) High parameter flow cytometry analysis of VP6:I-A^b+^ CD4^+^ T cells. t-SNE plot showing the distribution of cT_FH_, T_FH_1 and T_H_1 cells vis-a-vis their expression of CXCR6, PSGL-1 and BCL6, and histogram showing their relative expression of BCL6.

**Figure S5.**
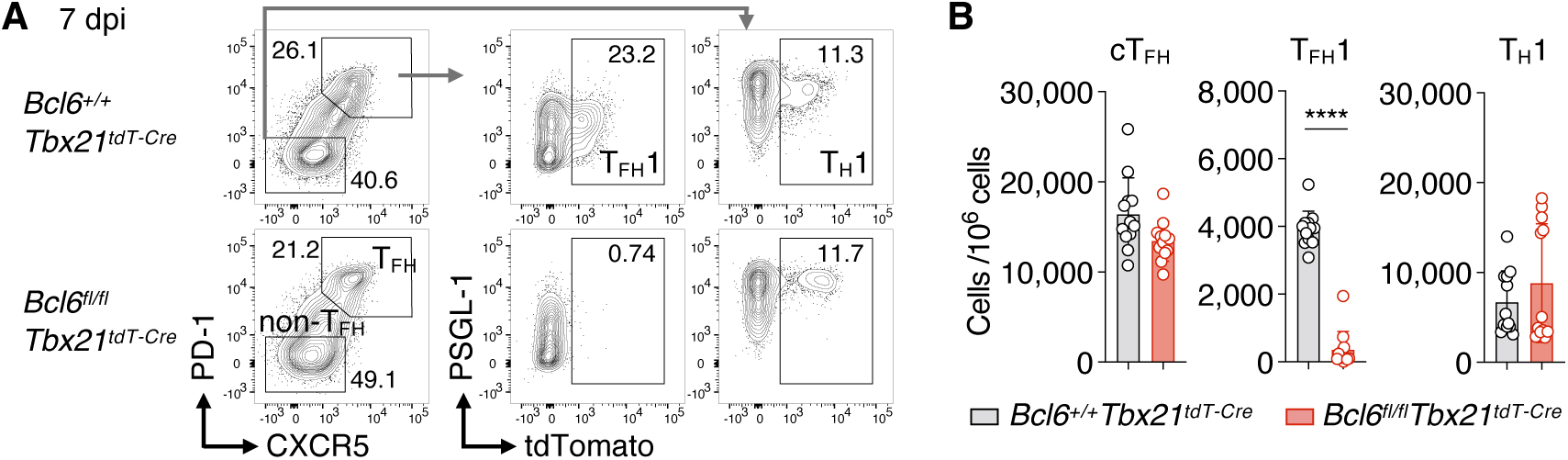
Analysis of mixed-BM chimeras deficient for T_FH_1 cells. (**A and B**) Flow cytometric analysis of PP cells in 80:20 mixed-BM chimeras employing *Tcrb^-/–^* BM and either *Bcl6^+/+^Tbx21^tdT-Cre^ or Bcl6^fl/fl^Tbx21^tdT-Cre^* BM as shown in Figure 5A at 7 dpi with RV. Representative plots showing the percentage of tdTomato^+^ cells among T_FH_ and non-T_FH_ cells (A) and graph showing the number of cT_FH_ (tdTomato^-^ T_FH_), T_FH_1 (tdTomato^+^ T_FH_) and T_H_1 (tdTomato^+^ non-T_FH_) cells (B). Data are pooled from 2 independent experiments and shown as mean ± SD; each symbol represents one mouse (n = 12); Mann-Whitney test.

**Figure S6.**
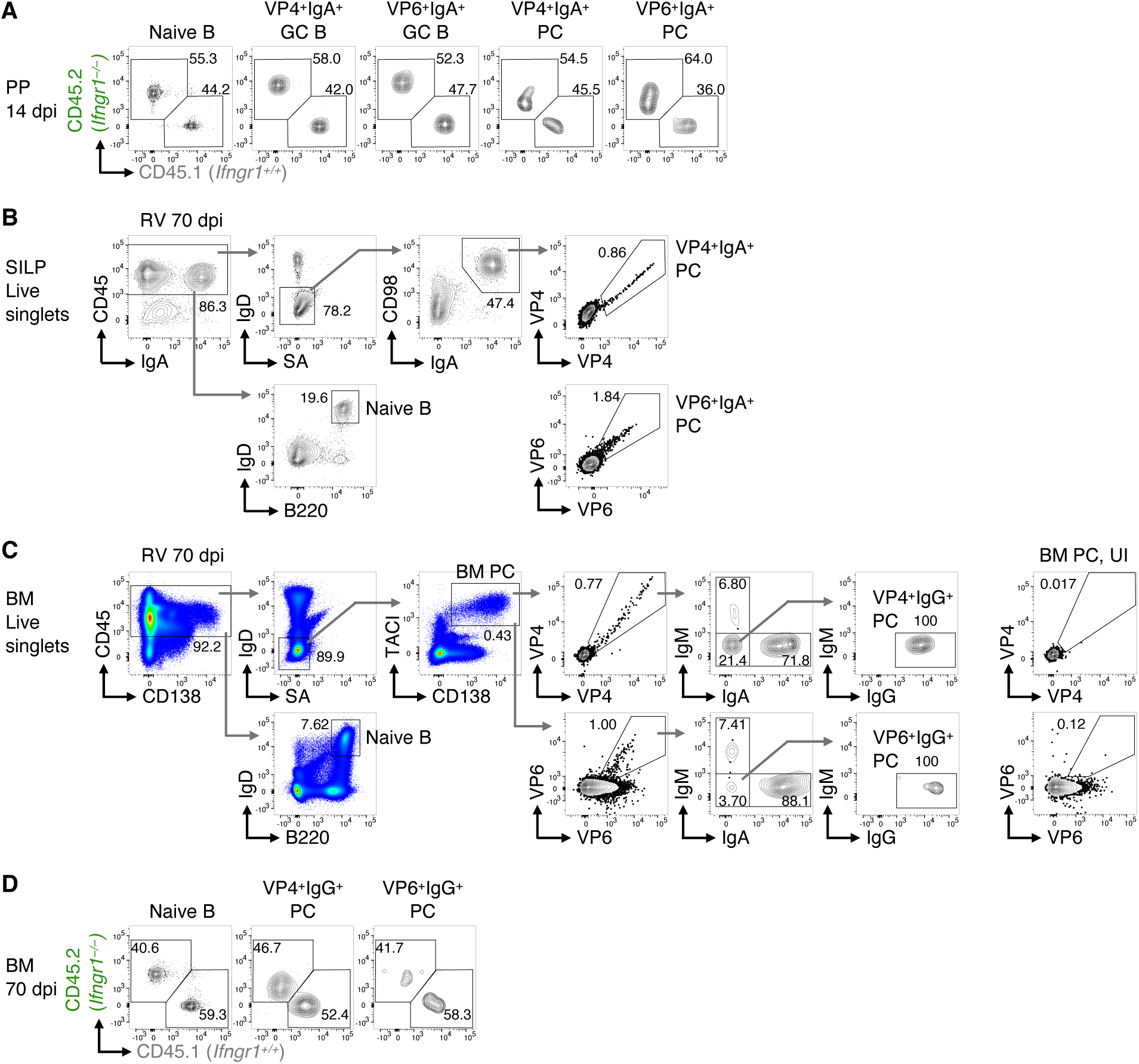
Flow cytometric analysis of 50:50 BM chimeras. (**A**) Flow cytometric analysis of 50:50 CD45.1-*Ifngr1*^+/+^ and CD45.2-*Ifngr1*^-/–^ mixed-BM chimeras. Representative flow cytometry plots showing the percentage of CD45.1^+^ and CD45.2^+^ cells in the PPs at 14 dpi. Quantification of the data is shown in Figure 6B. (**B**) Representative flow cytometry profiles showing the gating strategy of naive B cells and RV-specific IgA^+^ PCs in the SILP of mixed BM chimeras infected with RV. All Abs and Ag-tetramers (VP4 and VP6) were used for cell surface staining. (**C**) Representative flow cytometry profiles showing the gating strategy of naive B cells and RV-specific PCs in the BM of mixed BM chimeras uninfected (UI) or infected with RV. Abs to IgM, IgG, IgA and Ag-tetramers (VP4 and VP6) were used for intracellular staining. (**D**) Flow cytometric analysis of 50:50 CD45.1-*Ifngr1*^+/+^ and CD45.2-*Ifngr1*^-/–^ mixed-BM chimeras. Representative flow cytometry plots showing the percentage of CD45.1^+^ and CD45.2^+^ cells in the BM at 70 dpi. Quantification of the data is shown in Figure 6F.

**Figure S7.**
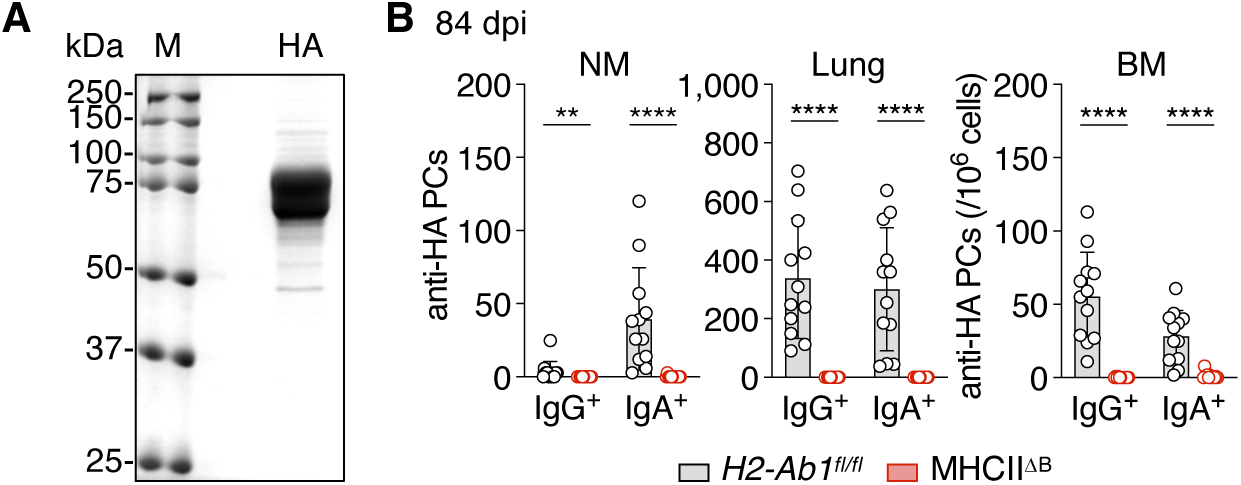
Analysis of PR8 IAV infections. (**A**) SDS-PAGE of purified recombinant PR8-HA. SDS-PAGE gels under reducing condition were stained with Coomassie brilliant blue. M: molecular weight marker. (**B**) Enumeration of HA-specific IgG^+^ and IgA^+^ PCs at 84 dpi with PR8 in *H2-Ab1^fl/fl^* and MHCII^ΔB^ mice. ELISPOT data are pooled from 2 independent experiments and shown as mean ± SD; each symbol represents one mouse (n = 12); Mann-Whitney test.

